# Proliferation control of kidney interstitial cells

**DOI:** 10.1101/2020.04.11.037259

**Authors:** Sarah S. McCarthy, Lindsey Gower, Michele Karolak, Alicia England, Thomas Carroll, Leif Oxburgh

**Author notes:** Author for correspondence:. Mailing address: The Rogosin Institute, 310 East 67th Street, New York, NY 10065.

## Abstract

Expansion of interstitial cells in the adult kidney is a hallmark of chronic disease, whereas their proliferation during fetal development is necessary for organ formation. An intriguing difference between adult and neonatal kidneys is that the neonatal kidney has the capacity to control interstitial cell proliferation when the target number has been reached. In this study, we define the consequences of inactivating the TGFβ/Smad response in the interstitial cell lineage. We find that pathway inactivation through loss of *Smad4* leads to over-proliferation of interstitial cells regionally in the kidney medulla. Genetic and molecular interaction studies showed that Smad3/4 participates in the Wnt/β-catenin signaling pathway, which is responsible for promoting proliferation of interstitial cells. Specifically, *Smad4* is required for the expression of the Wnt feedback inhibitor *Apcdd1*, and based on these findings we propose a model for interstitial cell proliferation control in which the Wnt/β-catenin proliferative signal is attenuated by TGFβ/Smad signaling to ensure that proliferation ceases when the target number of interstitial cells has been reached in the neonatal medulla.

**Summary statement:** This study describes a novel function for TGFβ signaling in the developing renal interstitium. Mice with Foxd1-Cre-mediated deletion of Smad4 have interstitial expansion and activated Wnt signaling.

## INTRODUCTION

Interstitial cells are any cells located between the functional cells of a tissue. In the kidney, cells of the nephron and blood vessel are considered the functional components and the interstitial cell population is made up largely of PDGFRβ-expressing fibroblasts. These cells derive from the *Foxd1* expressing progenitor, which also gives rise to mesangial cells, the specialized pericytes of the glomerulus. Expansion of interstitial cells in the adult kidney is a hallmark of chronic disease, with unopposed interstitial cell proliferation leading to progressive scarring (fibrosis) and concomitant loss of parenchyme (Humphreys et al., 2010). In contrast, proliferation of interstitial cells plays an essential role in fetal development and somatic growth of the kidney (Boivin and Bridgewater, 2018; Das et al., 2013; Fetting et al., 2014; Hatini et al., 1996). An intriguing difference between adult and neonatal kidneys is that the neonatal kidney has the capacity to control interstitial cell proliferation when the target number has been reached, and our study aims to characterize basic mechanisms of this proliferation control.

Gene inactivation and experimental therapeutics targeting the TGFβ signaling pathway have been used to reduce fibrosis in kidney injury models (Inazaki et al., 2004; Morishita et al., 2014), indicating that the pathway is a significant driver of interstitial expansion. We therefore hypothesized that the TGFβ pathway may control interstitial cell proliferation in the neonatal kidney. Although TGFβ signaling has been a focus of research on homeostasis of adult interstitial cells, little is known about its function in differentiation of the renal interstitium. TGFβ signaling is essential for development of the kidney (Dudley et al., 1995; Ikeya et al., 2010; Oxburgh et al., 2004), and controls formation of both the collecting ducts and nephrons (Brown et al., 2013; Hartwig et al., 2008). Recessive mutations in genes within this signaling pathway have been identified as monogenic causes of Congenital Anomalies of the Kidney and Urinary Tract (CAKUT), indicating that TGFβ pathway dysregulation may be an important factor in neonatal kidney disease. An important example is the identification of mutations in bone morphogenetic protein 4 (BMP4) in patients with renal hypodysplasia (Kohl et al., 2014; Weber et al., 2008).

TGFβ superfamily ligands signal through two distinct intracellular pathways: Smad, which is initiated by phosphorylation of receptor-associated Smad (R-Smad) transcription factors and mitogen-associated protein kinase (MAPK) which is initiated by the TGFβ-associated kinase TAK1 (MAP3K7). Kidneys of mice with conditional inactivation of *Map3k7* in the *Foxd1* lineage develop spontaneous neonatal mesangiosclerosis (Karolak et al., 2018), suggesting that MAPK is selectively required for mesangial differentiation.

The current study addresses the role of Smad pathway signaling in the *Foxd1* lineage. Phosphorylated R-Smads associate with the common mediator Smad (Smad4) to accumulate in the nucleus. R-Smads and Smad4 bind DNA and interact with a variety of other transcription factors, facilitating highly context-dependent responses. Because Smad4 is unique and essential for Smad-mediated responses, we selected it as a tractable node in the pathway for conditional gene inactivation. We report that *Smad4* controls proliferation within the interstitium of the neonatal kidney by attenuating Wnt signaling.

## RESULTS

### Loss of *Smad4* causes expansion of the renal interstitium

To understand the role of TGFβ/BMP in the developing renal interstitium, *Smad4* was inactivated in interstitial cell progenitors using *Foxd1*^*Cre*^. To sensitize the strain for cre-mediated recombination, *Foxd1*^*Cre*^ was combined with one null allele for *Smad4* and one loxP-flanked allele (Chu et al., 2004). *Foxd1*^*+/Cre*^;*Smad4*^*-/loxp*^ is referred to as *Smad4*^*IC*^ and the *Foxd1*^*+/Cre*^;*Smad4*^*+/loxp*^ control is referred to as *Smad4*^*con*^. To evaluate the efficiency of cre recombination, we first compared *Smad4* transcript levels, and found that they are reduced by approximately 60% in *Smad4*^*IC*^ kidneys versus *Smad4*^*con*^ (Fig. 1A). To measure recombination efficiency specifically in the *Foxd1* lineage, we selected cells expressing the interstitial cell surface marker PDGFR*α* (Fig. S1A,B), and single-cell genotyped them (Fig. S1C,D). Figure 1B summarizes the frequency of recombination of interstitial cells in the *Smad4*^*IC*^ genetic model; 87.1% of cells were recombined at the *Smad4* locus and thus null for *Smad4*.

**Figure 1.**
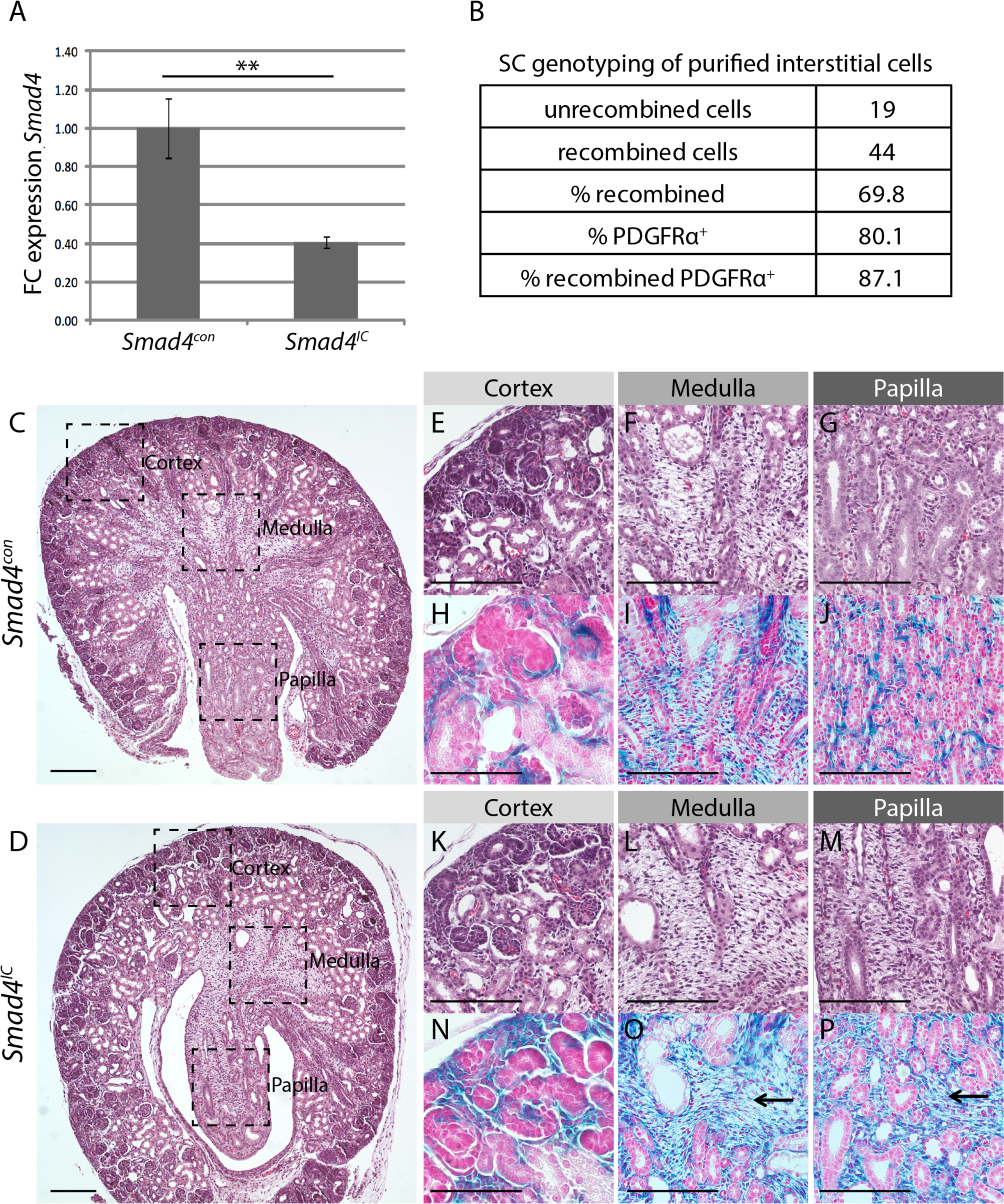
Single cell recombination analysis and histology of *Smad4*^*con*^ and Smad4^*IC*^ postnatal kidneys. (A) *Smad4* transcript levels in whole kidneys isolated from *Smad4*^*con*^ and *Smad4*^*IC*^ P0 mice (**=p<0.01; n=6). FC = fold change; transcript levels normalized to *Smad4*^*con*^ ± SEM from three separate experiments are graphed. (B) Summary of recombination frequency in purified cortical interstitial cells. (C, D) Transverse kidney sections with boxes showing regions of cortex, medulla and papilla shown at higher magnification in panels E-P. (E, F, G) Representative H&E fields of *Smad4*^*con*^ kidneys. (H, I, J) Representative fields of X-Gal stained sections from kidneys of *Smad4*^*con*^ on the R26R background. (K, L, M) Representative H&E fields of *Smad4*^*IC*^ kidneys. (H, I, J) Representative fields of X-Gal stained sections from kidneys of *Smad*^*IC*^ on the R26R background. Arrows denote X-Gal-positive regions of expanded stroma. Scale bars: 200µm in C, D; 100µm in E-P.

To determine if loss of *Smad4* in the *Foxd1*-expressing interstitial cell progenitor affects lineage commitment, we introduced the *Rosa26R* background. The localization of *Foxd1* lineage cells in the *Smad4*^*IC*^;*Rosa26R* strain is indistinguishable from *Smad4*^*con*^;*Rosa26R*, and thus we conclude that lineage commitment is unperturbed (Fig. S2). Transverse sections through the kidney reveal extensive stroma in the medulla and papilla of *Smad4*^*IC*^, and a paucity of epithelial structures in both of these zones of the mutant (Fig. 1C,D). The R26R reporter serves as a helpful marker that can be used in parallel with histological analysis to understand the abundance of *Foxd1* lineage cells. Comparison of *Smad4*^*con*^ (Fig. 1 E-J) and *Smad4*^*IC*^ (Fig. 1K-P) revealed a marked expansion of *Foxd1*-lineage interstitium which was regionalized in the tissue; little difference was noted in cortical interstitium (Fig. 1H,N), while pronounced pockets of stroma were seen in the medulla and papilla (Fig. 1I,J,O,P). We conclude that *Smad4* is required in the *Foxd1* lineage for appropriate formation of the renal interstitium.

### Smad4 is required for appropriate differentiation of interstitial cells

During renal development, interstitial cells transition from a FOXD1/PDGFRα-positive state in the cortex to a PDGFRβ/α-SMA-positive state in the medulla. To determine if the stromal expansion observed in *Smad4*^*IC*^ mice is associated with impaired interstitial cell differentiation, expression of the medullary interstitial markers PDGFRβ and α-SMA were examined in kidneys from embryonic and postnatal mice. Though comparable to control at E17.5 (Fig. S3), differences in α-SMA and PDGFRβ expression are observed in *Smad4*^*IC*^ mouse kidneys by P0. *Smad4*^*IC*^ mice display reduced medullary α-SMA expression compared to control (Fig. 2A,B). In contrast, PDGFRβ expression is increased in the inner and outer medulla of *Smad4*^*IC*^ mice compared to control (Fig. 2C,D). Confirming these findings, immunoblots of lysates from whole kidneys (Fig. 2E) showed a significant increase of PDGFRβ (Fig. 2F) and decrease of α-SMA (Fig. 2G) in *Smad4*^*IC*^ mice. The lack of expression of α-SMA in the abundant interstitial cells of the medulla suggests that *Smad4* is required for their differentiation.

**Figure 2.**
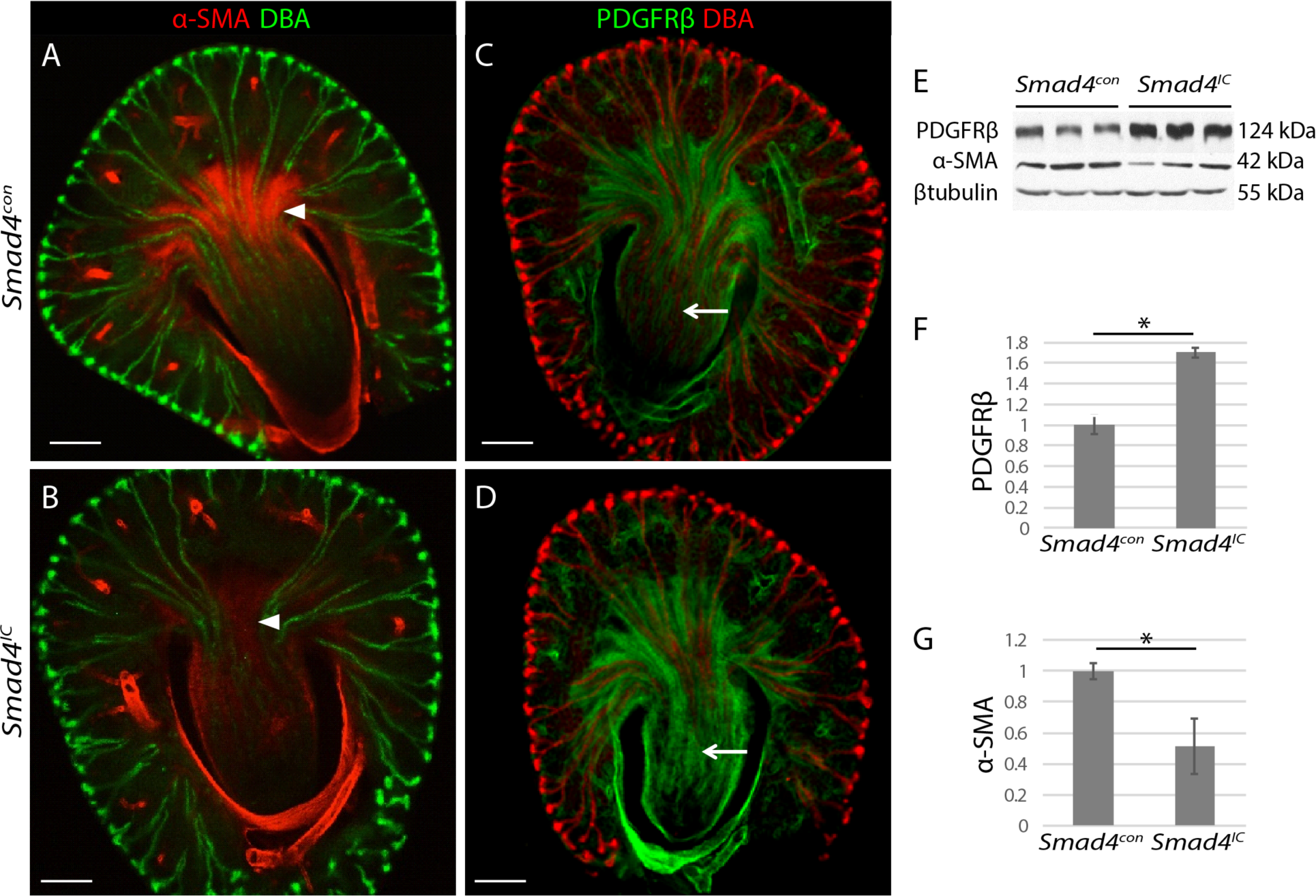
*Smad4* is required for differentiation of the renal interstitium. Whole mount immunofluorescence of kidney vibratome sections representative of 6 *Smad4*^*con*^ and 6 *Smad4*^*IC*^ kidneys stained with DBA and α-SMA (A, B) or PDGFRβ (C, D). Arrowheads mark comparable regions of medullary stroma and arrows indicate comparable regions of papillary stroma in *Smad4*^*con*^ and *Smad4*^*IC*^. (E) Immunoblots of whole kidney lysates from three *Smad4*^*con*^ and three Smad4^*IC*^ mice probed with α-SMA, PDGFRβ, and β-tubulin antibodies. (F) Quantification of band intensities for PDGFRβ. (G) Quantification of band intensities for *α*-SMA. Error bars are SEM and represent three independent experiments. *=p<0.05. Scale bars: 200µm.

### Loss of Smad4 from the Foxd1 lineage causes features of collecting duct compression and urine outflow occlusion

One possible explanation for the pockets of interstitial cells seen in the *Smad4*^*IC*^ medulla is that collecting duct (CD) organization is impaired, causing aberrant clustering of interstitial cells, and to determine if this was the case we compared *Smad4*^*IC*^ kidney tissue with controls using two strategies. Three dimensional modeling of whole kidneys stained with the CD marker TROMA1 showed that branching at embryonic day 14.5 is indistinguishable between *Smad4*^*con*^ and *Smad4*^*IC*^ mice (Fig. 3A-C, S4A,B). To analyze collecting ducts at P0 when the kidney is too large for modeling based on immunostaining, we sectioned through the longitudinal plane of the kidney perpendicular to the papilla to obtain transverse sections of the papilla where collecting ducts are closely bundled (Fig. 3D), and immunostained these sections. We found a reduced number of patent TROMA1-positive structures in the papilla of *Smad4*^*IC*^ mice compared to *Smad4*^*con*^ (Fig. 3E,F). High magnification images of the papilla revealed TROMA1-positive tubules with an atypical compressed morphology in *Smad4*^*IC*^ kidneys (Fig. 3G,H). When quantified, the compressed TROMA1-expressing CDs account for the reduced number of CDs in *Smad4*^*IC*^ mice (Fig. 3I). We hypothesize that these structures are remnants of functional CDs that are constricted by stromal expansion. CD compression is predicted to cause restricted urine outflow, and consistent with this, *Smad4*^*IC*^ mutant kidneys displayed nephron tubule expansion (Fig. 3J-L) and distended Bowman’s spaces (Fig. 3M-O). Increased mesangial α-SMA expression (Fig. 3P,Q, S4C,D) and mesangial area (Fig. 3R) indicate early glomerulosclerosis in *Smad4*^*IC*^ mice. *In vivo* 5-ethynyl-2’-deoxyuridine (EdU) incorporation revealed an increased proliferative index in mesangial cells of the mutant at P0 (Fig. S4E-G), indicating that this defect is mesangioproliferative. Numbers of glomeruli are comparable between *Smad4*^*con*^ and *Smad4*^*IC*^ (Fig. S4H), suggesting that the features of increased physiological load that we observed are not due to differences in nephron number. In summary, the histological features that we observed are consistent with CD compression causing increased luminal pressure in the nephron with associated sclerosis of the glomerulus.

**Figure 3.**
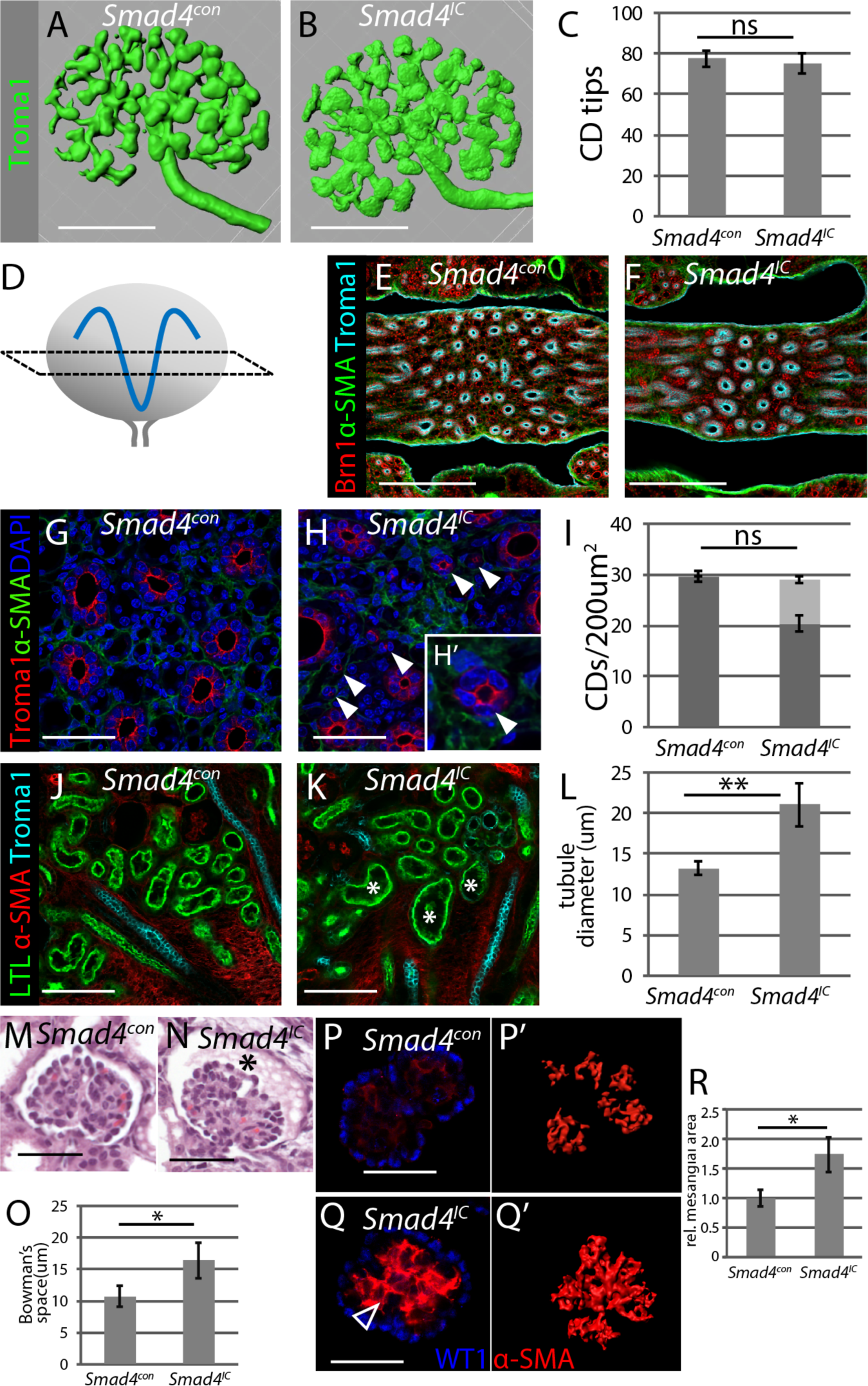
Interstitial expansion results in collecting duct constriction, tubular distension, and expansion of Bowman’s space. 3-D reconstruction of E14.5 kidneys from *Smad4*^*con*^ (A) and *Smad4*^*IC*^ (B) mice stained for Troma1. (C) Quantification of collecting duct tip number per E14.5 kidney (n=6). (D) Schematic of sectioning plane used for P0 collecting duct analysis. (E, F) P0 kidney vibratome sections stained with Brn1, α-SMA, Troma1. (G, H) P0 kidney vibratome sections stained with α-SMA, Troma1, DAPI. Arrowheads mark abnormally dimensioned Troma-1^+^ collecting ducts. (I) Quantification of collecting ducts per 200µm^2^ papilla (n=6). (J, K) P0 kidney vibratome sections stained with LTL, α-SMA, Troma1. Asterisks mark distended proximal tubules. (L) Quantification of LTL+ proximal tubule diameter (n=6). (M, N) Representative sections of glomeruli. Asterisk denotes distended Bowman’s space. (O) Quantification of Bowman’s space from 6 individual kidneys of each genotype. Only glomeruli that sectioned through the center were counted. (P, Q) Representative sections through P0 glomeruli co-stained with WT1 and *α*-SMA. Arrowhead marks increased mesangial α-SMA expression. (P’, Q’) Maximal intensity projections of Z-stacks through glomeruli immunostained with α-SMA. (R) Relative mesangial area normalized to *Smad4*^*con*^ (µm^2^; n=6) of glomeruli from *Smad4*^*con*^. *=p<0.05; **=p<0.01. Scale bars: 200µm in A-F; 50µm in G-K; 20µm in M-P.

### Loss of *Smad4* affects nuclear accumulation of Smad3 but not Smad1/5/8

To understand if loss of *Smad4* affects nuclear accumulation of R-Smads in regions of medullary interstitial cell expansion, protein localization of Smads was compared in *Smad4*^*con*^ and *Smad4*^*IC*^ mouse kidneys. To localize expression in interstitial cells, we co-stained with *α*SMA. Signal amplification was used to reliably detect *α*SMA in the *Smad4*^*IC*^ kidney. As anticipated, Smad4 was lost from the interstitium of *Smad4*^*IC*^ mice (Fig. 4 A,B). In addition, nuclear accumulation of Smad3 was strongly reduced in the interstitium of *Smad4*^*IC*^ kidneys compared to control (Fig. 4C,D), while nuclear pSmad1/5/8 levels were only slightly reduced (Fig. 4E,F). These results reveal that nuclear accumulation of Smad3, but not Smad1/5/8, is dependent on Smad4 in medullary interstitial cells and that the TGFβ signaling pathway is primarily affected by *Smad4* inactivation.

**Figure 4.**
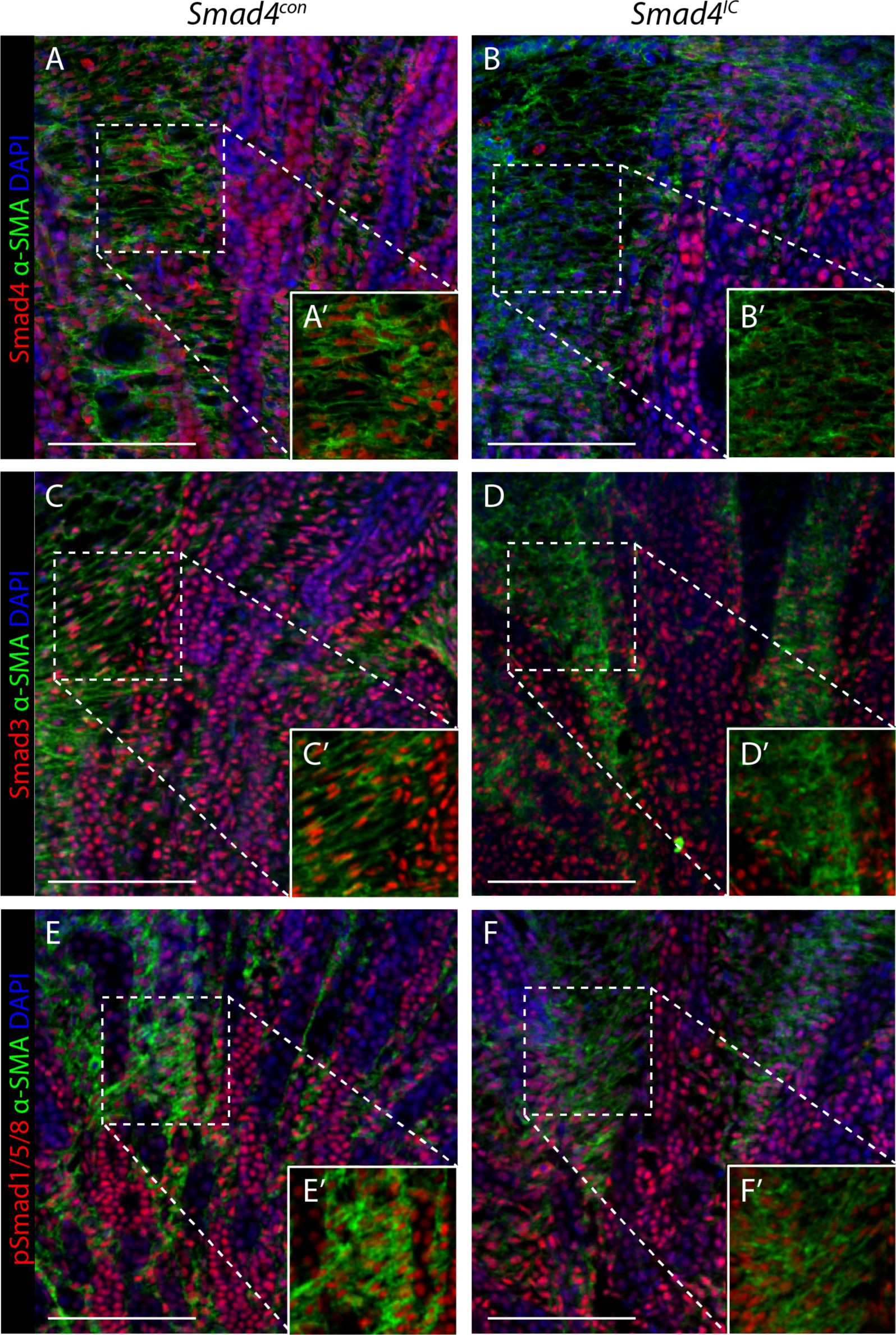
Nuclear Smad3, but not Smad1/5/8, is decreased in the interstitium of *Smad4*^*IC*^ mice. Immunofluorescence with Tyramide Signal Amplification (TSA) of P0 kidneys from *Smad4*^*con*^ (A, C, E) and *Smad4*^*IC*^ (B, D, F) mice stained with antibodies recognizing α-SMA (green), DAPI (blue) and Smad4 (A, B), Smad3 (C, D) or pSmad1/5/8 (E, F); n=6. Scale bars: 100µm.

### Interstitial expansion is due to increased proliferation

To determine whether aberrant proliferation is responsible for the interstitial expansion observed in *Smad4*^*IC*^ mice, we performed *in vivo* EdU incorporation at P0 (Fig. 5). To localize proliferating cells in the interstitium, we co-stained tissue with a cocktail of antibodies for *α*SMA and PDGFRβ. While EdU^+^ cycling interstitial cells were largely limited to the cortex and outer medulla in *Smad4*^*con*^ kidneys, they were abundant in the inner medulla of *Smad4*^*IC*^ mice (Fig. 5A,B). 3D modeling and quantification of comparable volumes confirmed a higher proliferation rate in the inner medullary interstitium (IMI) of *Smad4*^*IC*^ mice compared to *Smad4*^*con*^ (Fig. 5C-E). We conclude that there is a failure of interstitial cell growth restriction in the transition zone between outer and inner medulla.

**Figure 5.**
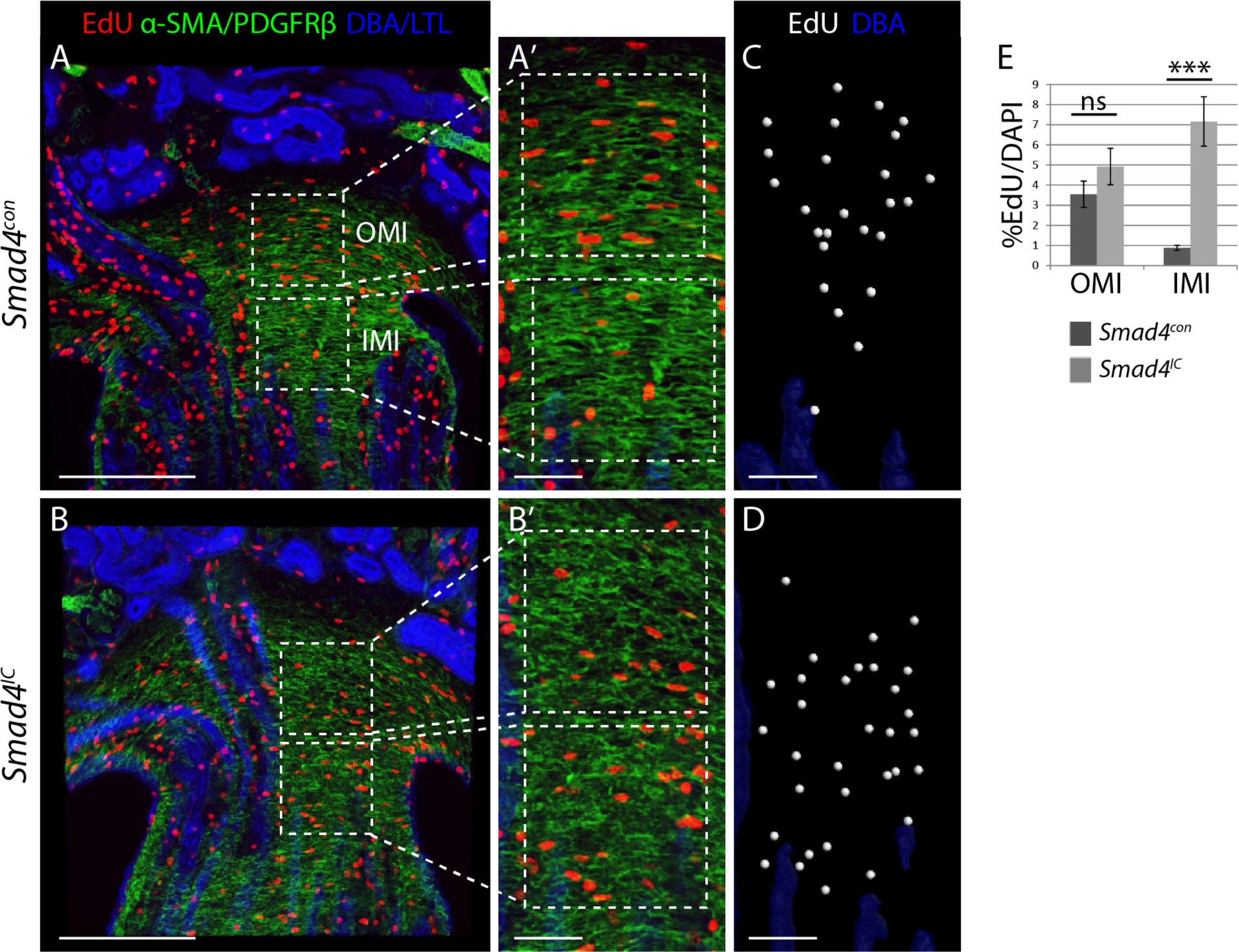
Loss of *Smad4* leads to increased interstitial proliferation. EdU labeling of kidney sections co-stained with collecting duct marker DBA, proximal tubule marker LTL, and α-SMA/PDGFRβ from *Smad4*^*con*^ (A, A’) and *Smad4*^*IC*^ (B, B’) mice. (C, D) Three-dimensional tissue analysis of EdU^+^ cells in hatched volumes of the outer medullary interstitium (OMI) and inner medullary interstitium (IMI) using Imaris software. (E) Quantification of 3-D analysis showing %EdU^+^/DAPI^+^ interstitial cells in the hatched volumes of OMI and IMI of *Smad4*^*con*^ and *Smad4*^*IC*^ mice; n=6. ns=p>0.05; ***=p<0.001. Scale bars: 200µm in A, B; 50µm in A’, B’, C, D.

### Interstitial expansion correlates with aberrant Wnt/β-catenin signaling

Several lines of evidence indicate that Wnt/β-catenin signaling is required for interstitial cell maintenance: loss of *Wnt7B* from the collecting duct epithelium or inactivation of β-catenin in interstitial cell precursors both result in medullary hypoplasia (Boivin et al., 2016; Yu et al., 2009). Thus, developmental genetic studies support a model in which collecting duct-derived Wnt drives medullary interstitial cell proliferation through a β-catenin dependent signaling mechanism. We were therefore curious to understand if Wnt/β-catenin signaling was perturbed in the *Smad4*^*IC*^ kidney. Expression of feedback inhibitors is a sensitive read-out of Wnt/β-catenin signaling. Analysis of transcriptome data suggested that the Wnt inhibitor *Apcdd1* is expressed in medullary interstitial cells (England et al., 2020), and we compared its expression level in *Smad4*^*con*^ and *Smad4*^*IC*^ kidneys. In situ hybridization revealed strong and specific *Apcdd1* expression in the medulla, which was drastically reduced in *Smad4*^*IC*^ (Fig. 6A,B). Reduction in activation of feedback inhibitors can be interpreted as evidence of reduced Wnt/β-catenin signaling, which would be unanticipated considering the proliferative phenotype of *Smad4*^*IC*^. We therefore evaluated expression of a panel of other Wnt/β-catenin targets. LEF1 was elevated in the medullary interstitium of *Smad4*^*IC*^ kidneys (Fig. 6C-E), along with the cell cycle regulator CCND1 (Fig. 6F-H), suggesting an increase in Wnt/β-catenin signaling and proliferation relative to *Smad4*^*con*^. We also found increased expression of the Wnt responsive cell cycle regulator CDKN1C (P57KIP2) (Fig. 6I-K), which is required for renal medulla formation (Yu et al., 2009; Zhang et al., 1997).

**Figure 6.**
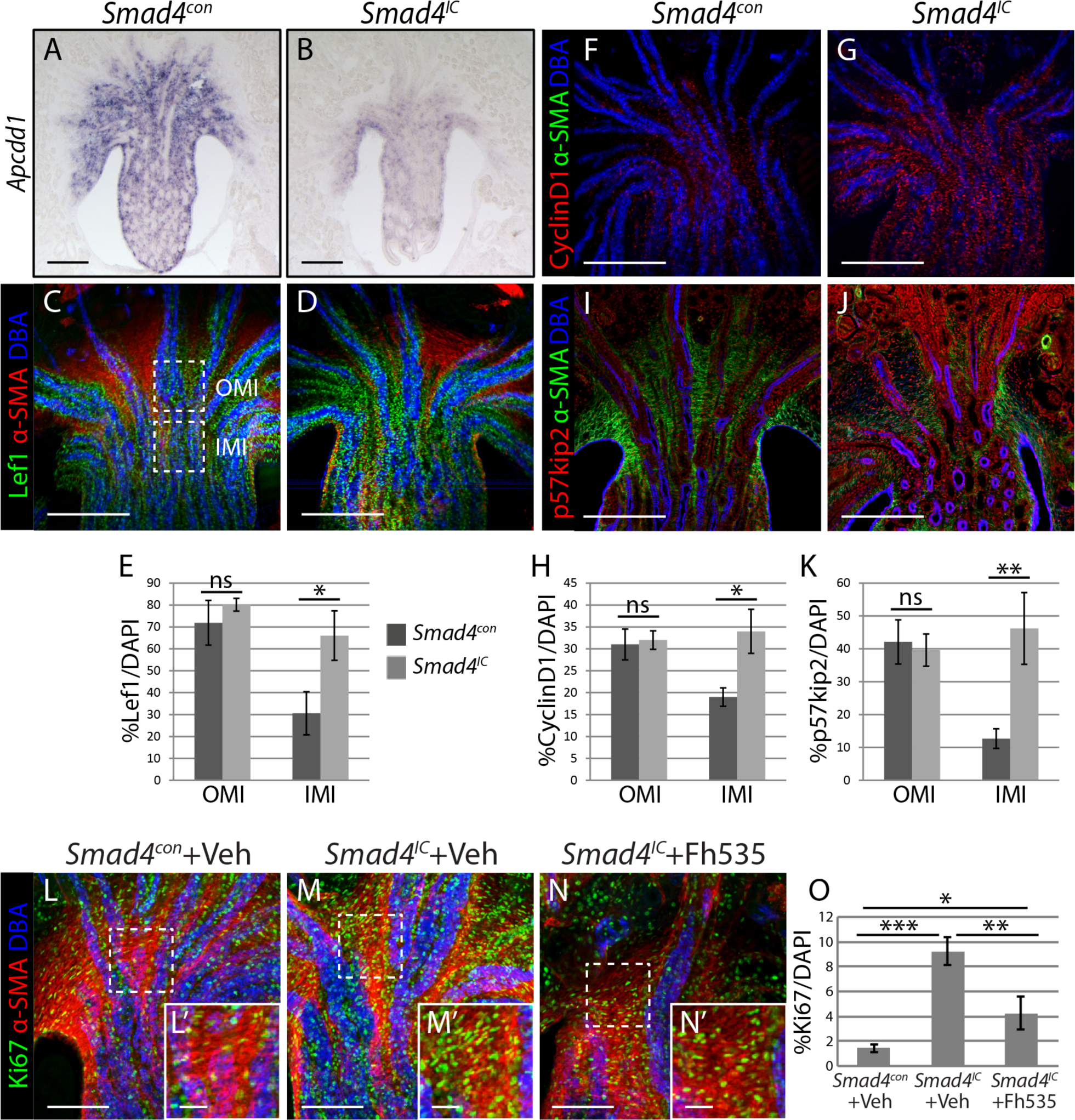
Increased proliferation is associated with aberrant Wnt signaling. (A, B) *Apcdd1 in situ* hybridization on P0 kidneys from *Smad4*^*con*^ (A) and *Smad4*^*IC*^ (B) mice. Immunofluorescent staining of vibratome sections of P0 kidneys from *Smad4*^*con*^ (C, F, I) and *Smad4*^*IC*^ (D, G, J) mice stained with antibodies recognizing α-SMA and DBA combined with either LEF1 (C, D), CyclinD1 (F, G) or p57kip2 (I, J). (E-K) Quantification of numbers of stained cells in the outer medullary interstitium (OMI) and inner medullary interstitium (IMI) of immunostained sections. Examples of regions selected for quantification are shown in panel C. (E) Quantification of LEF1^+^/DAPI^+^ (n=6), (H) Quantification of CyclinD1^+^/DAPI^+^ (n=6) (K) Quantification of p57kip2^+^/DAPI^+^ (n=6). Immunofluorescent staining of P0 kidneys from mice treated with either vehicle (L, M) or 20mg/kg FH535 (N). *Smad4*^*con*^ (L) and *Smad4*^*IC*^ (M, N) mice stained with Ki67, α-SMA, and DBA. (O) Quantification of Ki67^+^/DAPI^+^ cells in the interstitium of kidneys from treated mice (n=6). Examples of *α*-SMA-expressing regions for quantification are shown with boxes and magnified in L’-N’. ns=p>0.05; *=p<0.05; **=p<0.01; ***=p<0.001. Scale bars: 200µm in A-J; 100µm in L-N; 20µm in L’-N’.

In aggregate, these results suggest that *Smad4* restricts Wnt/β-catenin signaling through activation of Wnt feedback inhibition, notably through *Apcdd1*. In this scenario, loss of *Smad4* would lead to unopposed Wnt/β-catenin signaling resulting in sustained expression of cell cycle regulators and excessive proliferation. To test the hypothesis that Wnt/β-catenin is driving the excessive interstitial proliferation in *Smad4*^*IC*^ kidney, litters of neonatal mice were treated with the β-catenin transcription inhibitor FH535 in an attempt to rescue the mutant phenotype (Fig. 6L-N). Proliferation analysis 24 hours after inhibitor treatment revealed a reduction compared to vehicle (Fig. 6O), confirming that Wnt/β-catenin drives the interstitial cell proliferation caused by loss of *Smad4*.

### Interactions between Smad4 and Wnt pathway components at the cellular level

Smads modulate the Wnt signaling pathway in a number of developmental and disease contexts (Jia et al., 2014; Li et al., 2016; Lim and Hoffmann, 2006; Salazar et al., 2013; Stevens et al., 2017; Webber et al., 2016), and to establish a cellular model with which we could study pathway interactions in the interstitium we generated a primary cell line. A method was developed to purify primary renal interstitial cells (PRIC) from wild type Swiss Webster neonates (Fig. S5). Purified cells were transduced with a lentivirus expressing a temperature sensitive mutant of SV40T that is stable at 33°C and degraded at 37°C (Loeber et al., 1989; Paucha et al., 1986). Clonal cell lines propagated at 33°C were isolated from transduced primary interstitial cells. Clone 3-1 was selected for further characterization based on its temperature-sensitive loss of SV40T (Fig. S6A-D), transcriptional profile suggesting that it represents a medullary interstitial cell (England et al., 2020)(Fig. S6E), spindle-like morphology (Fig. 7A), and expression of the interstitial markers PDGFRβ (Fig. 7B), α-SMA (Fig. 7C), MEIS1, fibronectin and vimentin (Fig. S6F-H). To determine if clone 3-1 recapitulates the proliferative response to Wnt/β-catenin stimulation predicted from genetic studies, cells were treated with the Wnt/β-catenin agonist CHIR. A time-dependent increase in the Wnt pathway reporter *Lef1* was seen following treatment, verifying that clone 3-1 activates Wnt/β-catenin signaling in response to CHIR (Fig. S6I). EdU incorporation reveals that CHIR treatment does indeed promote proliferation of 3-1 interstitial cells (Fig. 7D), and abrogation of this effect by treatment with the β-catenin/TCF transcriptional inhibitor FH535 demonstrated that the effect is mediated through the β-catenin pathway (Fig. 7E). To determine the expression of TGFβ pathway components in 3-1 cells, we confirmed nuclear translocation of Smads in response to stimulation with BMP4 (Fig. S6J) or TGFβ1 (Fig. S6K). These features validate clone 3-1 as an in vitro model to investigate interactions between Smad4 and the Wnt pathway in interstitial cells and expression of key pathway components is summarized in Figure 7F.

**Figure 7.**
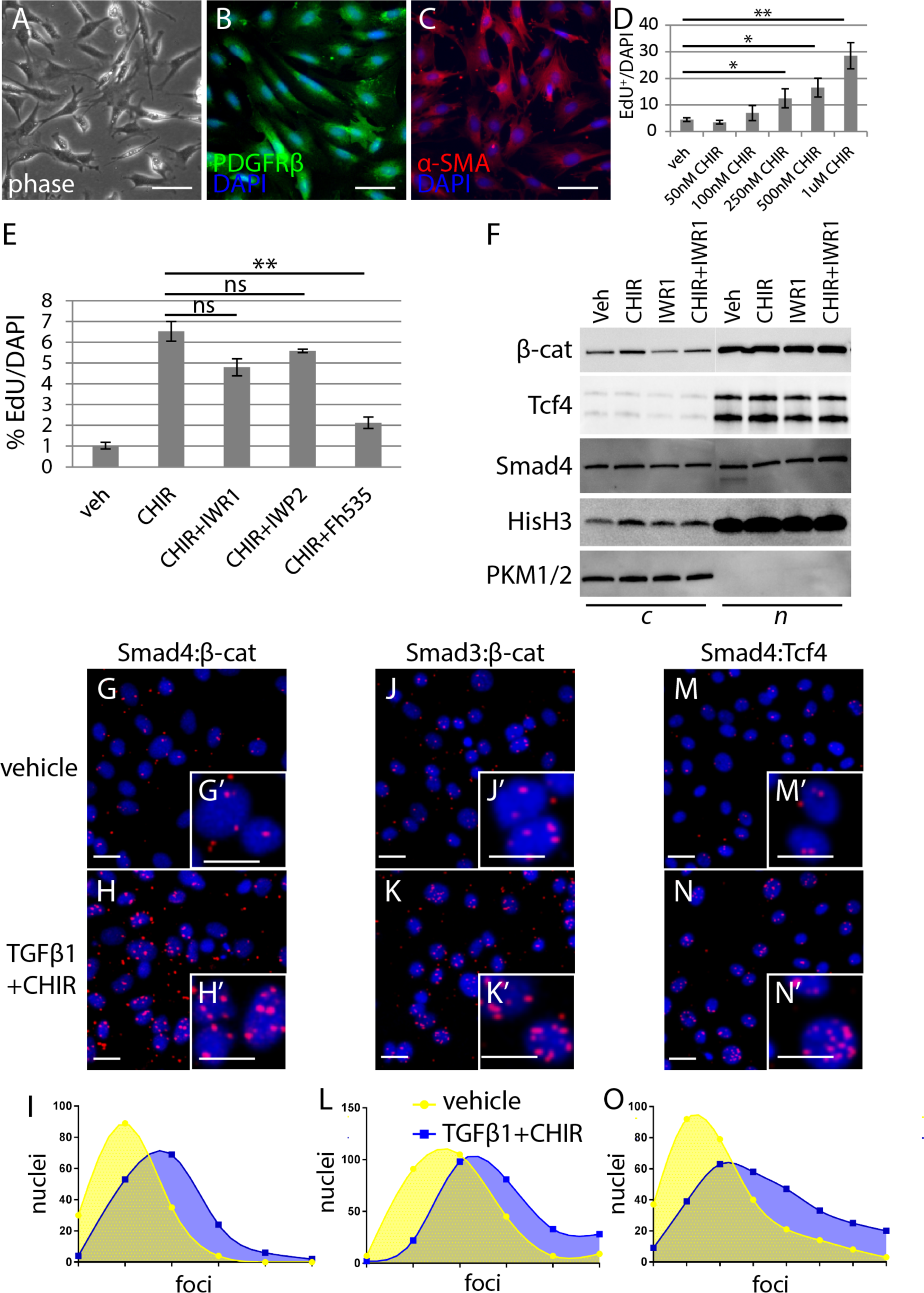
Smads colocalize with Wnt pathway components. (A) 3-1 interstitial cells imaged with phase contrast and immunostaining for PDGFRβ (B) or α-SMA (C). (D) Quantification of proliferative response (EdU^+^/DAPI^+^ cells) to increasing concentrations of CHIR. (E) Quantification of proliferation (%EdU/DAPI) in response to treatment with vehicle, 1µM CHIR alone, or co-treatment with 10µM IWR1, 5µM IWP2, or 15µM FH535 inhibitors. Error bars are SEM and represent three independent experiments. (F) Immunoblot for β-catenin, TCF4, and Smad4 in subcellular fractions of clone 3-1 treated with vehicle, CHIR, IWR1 or CHIR plus IWR1. HisH3 and PKM antibodies mark the nuclear (*n*) and cytosolic (*c*) fractions, respectively. (G-O) Fluorescence microscopy images and quantification of proximity ligation assay nuclear foci in cells probed with antibodies recognizing either Smad4+active β-catenin (G-I), Smad3+active β-catenin (J-L), or Smad4+TCF4 (M-O). Cells were treated with vehicle (G, J, M) or 5ng/ml TGFβ1 and 1µM CHIR (H, K, N). The graphs in I, L, O show counts of nuclear foci in vehicle treated cells (yellow) versus counts in TGFβ1/CHIR treated cells (blue). ns=p>0.05; *=p<0.05; **=p<0.01; ***=p<0.001. Scale bars: 10µm in A-C, G-N; 5µm in G’-N’.

To determine if there is a basis for participation of Smad4 in the β-catenin transcriptional complex, we used proximity ligation assay (PLA) to co-localize Smad4, Smad3 β-catenin, and the TCF4 transcriptional co-factor in 3-1 cells stimulated with CHIR and TGFβ1. In this assay, nuclear foci appear when antibodies bound to specific proteins are sufficiently close (0-40 nm) to prime an isothermal PCR reaction with ensuing fluorophore labeling. 3-1 cells co-stimulated with CHIR and TGFβ1 and probed with antibodies against Smad4 and β-catenin showed an increase in nuclear foci compared to vehicle-treated cells, indicating that they colocalize in the nucleus following stimulation (Fig. 7G-I). Similar results were observed when stimulated 3-1 cells were probed with Smad3:β-catenin (Fig. 7J-L) or Smad4:TCF4 (Fig. 7M-O) antibody combinations. These data show that Smad4, Smad3, β-catenin, and TCF4 all colocalize in close proximity to each other in the nucleus following co-stimulation of Wnt/β-catenin and TGFβ/Smad pathways, suggesting that Smad4 and Smad3 participate in the β-catenin/TCF4 transcriptional complex in interstitial cells.

### Smad4 drives expression of Apcdd1

Data to this point indicate that *Smad4* is required for Wnt/β-catenin feedback inhibition in interstitial cells. To test if *Smad4* is required for the expression of the Wnt/β-catenin antagonist *Apcdd1*, we generated a series of cell lines using the same approach as described for the 3-1 line (Fig. S5) from primary renal interstitial cells harvested from neonates of the *Hprt*^*CreERT2*^;*Smad4;R26R* genotype. The *Hprt* gene is ubiquitously expressed and because the *CreERT2* cassette is recombined into this locus, *Smad4* can be deleted by transient tamoxifen treatment. *Hprt* is located on the X chromosome, and therefore only male mice of *Hprt*^*CreERT2*^;*Smad4;R26R* genotype were used for cell line derivation. The bulk population from which clonal cell lines were derived showed dose-dependent activation of R26R reporter and loss of *Smad4* following tamoxifen treatment (Fig. S7A, B). Clones 50, 80, 82 and 87 were chosen for further analysis based on their recombination potential (Fig. S7C), significant tamoxifen-induced decrease in Smad4 protein (Fig. 8A,B), and transcriptional profile of interstitial markers (Fig. 8C). A decrease in *Apcdd1* transcript levels was observed in all four clones following tamoxifen treatment (Fig. 8D), demonstrating that Smad4 is required for the expression of the Wnt/β-catenin feedback inhibitor *Apcdd1*. Based on these findings we propose a mechanism in which Smad4/3 is required for activation of *Apcdd1*, which suppresses Wnt/β-catenin, thus limiting proliferation (Fig. 8E).

**Figure 8.**
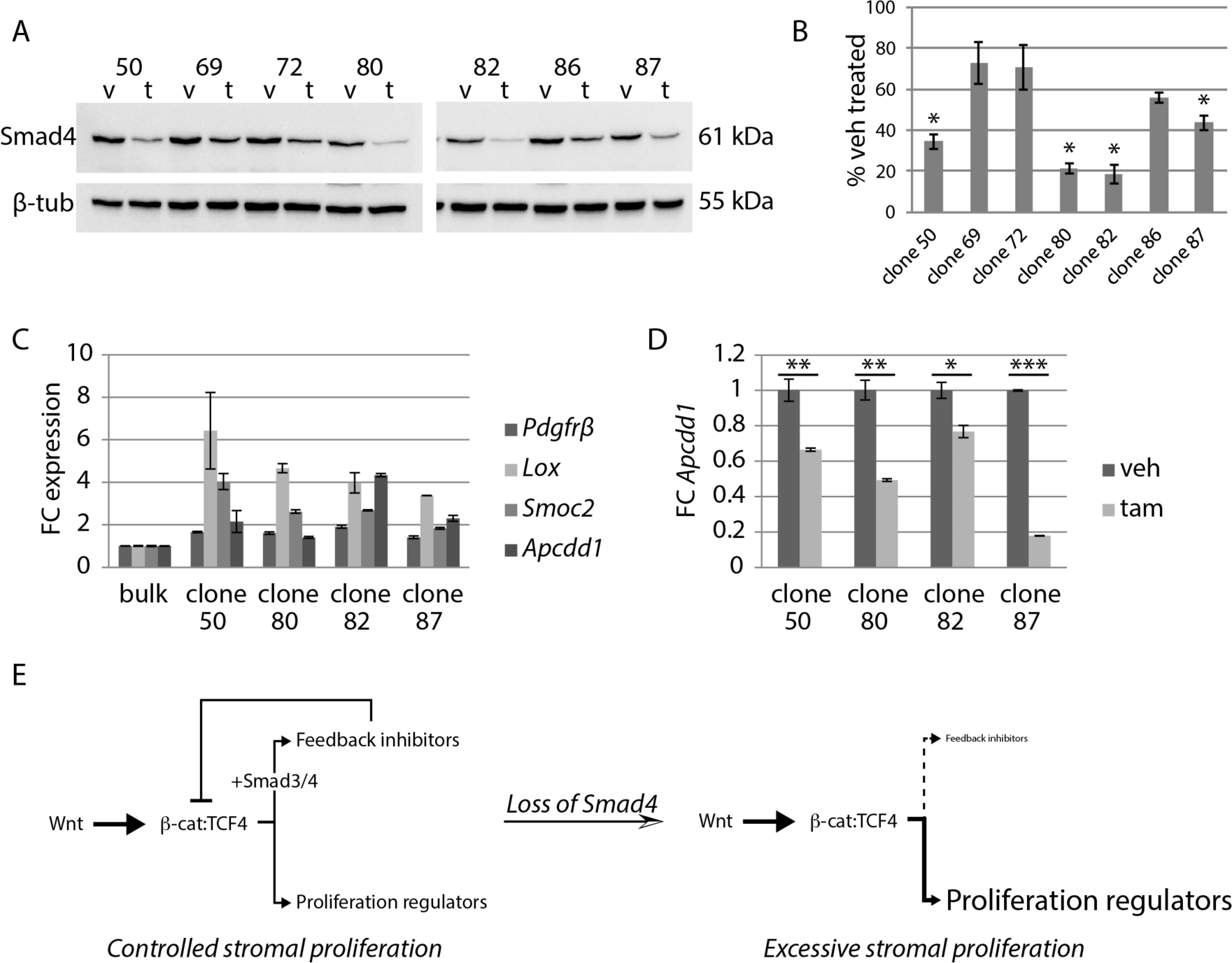
Smad4 drives expression of *Apcdd1*. (A) Smad4 immunoblot and band intensity quantification (B) of *Hprt*^*Cre*^;*Smad4;R26R* clones treated with vehicle or tamoxifen (1µg/ml). Percent vehicle treated is graphed and error bars (SEM) represent three independent experiments (*p<0.05 for clones 50, 80, 82, 87). (C) Fold change expression of interstitial cell markers *Pdgfrβ, Lox, Smoc2* and *Apcdd1* in *Hprt*^*Cre*^;*Smad4;R26R* clonal cell lines relative to the bulk population from which they were derived. (D) Fold change expression of *Apcdd1* in tamoxifen-treated *Hprt*^*Cre*^;*Smad4;R26R* clones relative to vehicle treated for clones 50, 80, 82, 87. (E) Schematic diagram representing the model for Smad-mediated control of Wnt-induced proliferation in interstitial cells based on findings in this study. *=p<0.05; **=p<0.01; ***=p<0.001.

## DISCUSSION

Considering the central role that *Smad4* plays in TGFβ superfamily signaling, it is not surprising that the *Foxd1*^*IC*^ strain displays a profound phenotype already in the neonate. TGFβ superfamily signaling in kidney fibroblasts has mainly focused on the study of TGFβ ligands, because foundational work showed that they promote proliferation (Roberts et al., 1985), extracellular matrix deposition (Edwards et al., 1987; Ignotz and Massagué, 1986), and myofibroblast transition (Rønnov-Jessen and Petersen, 1993). In previous work, we explored the consequences of inactivating the TGFβ/MAPK pathway in interstitial cells by inactivating TGFβ-associated kinase 1 (*Map3k7*) using *Foxd1*^*+/cre*^ (Karolak et al., 2018). In this study, we define the consequences of inactivating the TGFβ/Smad response by inactivating *Smad4*.

TGFβ promotes phosphorylation of the R-Smads 1, 2, 3, and 5 (Ramachandran et al., 2018; Zhang et al., 1996), and therefore one tractable strategy for studying Smad responses is to inactivate *Smad4*, which is required for nuclear retention and transcriptional activity of activated R-Smads (Lagna et al., 1996; Schmierer and Hill, 2005). Unexpectedly, we find that Smad4 inactivation eliminates nuclear accumulation of the Smad3 transcription factor, while only slightly reducing nuclear accumulation of the Smad1/5 transcription factors. This is in line with the Smad transcriptional response to TGFβ1, which is estimated to be comprised of approximately 75% Smad2/3 transcriptional response and 25% Smad1/5 transcriptional response (Ramachandran et al., 2018). The finding that Smad1/5 nuclear accumulation is not lost indicates that the BMP R-Smad response is only marginally affected by *Smad4* inactivation. Thus, from the perspective of Smad activation, our genetic model reflects the effects of eliminating TGFβ rather than BMP signaling.

The number of interstitial cells in the kidney must be carefully balanced both during development and in the adult. In development, interstitial cells provide essential signals that guide differentiation of the surrounding epithelia (Das et al., 2013; Fetting et al., 2014). In the adult, interstitial cells provide essential endocrine functions such as erythropoietin production (Kobayashi et al., 2016), but their uncontrolled expansion is the basis for organ fibrosis in which functional tissue is marginalized (Humphreys et al., 2010). Our observation that reducing TGFβ/Smad signaling in the interstitial cell lineage leads to increased proliferation in the neonate agrees with findings from studies of multiple cell lines showing that growth is inhibited by TGFβ (Roberts et al., 1985). An interesting feature of our study is that the effect is limited to a particular zone of interstitial cells in the medulla. Genetic studies have suggested that formation of the medullary interstitium requires Wnt/β-catenin signaling and our work indicates that TGFβ/Smad signaling inhibits this proliferative stimulus in a regional manner to ensure appropriate structural differentiation of the medulla. Interestingly, the effect of knocking out the Wnt/β-catenin pathway in the interstitial lineage is loss of the kidney medulla, but the cortex of these kidneys is preserved (Boivin and Bridgewater, 2018; Yu et al., 2009), suggesting that specificity of Wnt/β-catenin for medullary interstitium. Similarly, our inactivation study of *Smad4* reveals overproliferation of the medullary interstitium, indicating that this Wnt/β-catenin - TGFβ/Smad circuit for growth control is specific to this region.

Our findings support a model in which Smad4 and Smad3 interact with Wnt/β-catenin to modify signaling output. Proximity ligation (PLA) assays show that β-catenin, TCF4, Smad3, and Smad4 are in close proximity, suggesting physical interaction. Over-expression studies have previously demonstrated that physical interaction of Smad4 in the β-catenin transcriptional complex alters transcriptional output of Wnt signaling in the xenopus animal cap (Nishita et al., 2000), confirming the possibility that the effect of Smad3:Smad4 in the β-catenin transcriptional complex may be based on a direct physical interaction.

Loss of *Smad4* causes differential effects on Wnt/β-catenin targets, with a loss of expression of the Wnt feedback inhibitor *Apcdd1* and increased LEF1 and p57kip2. This supports a model in which loss of feedback inhibition causes increased Wnt/β-catenin signaling resulting in increased proliferation, which is represented in the model shown in Fig. 8E. Based on our findings, we propose that the loss of feedback inhibition is a primary event in the deregulation of interstitial cell proliferation. Multiple Wnts including *Wnt4* and *Wnt7b* are expressed in the medullary interstitium (Yu et al., 2009), facilitating combinatorial effects. TGFβ sources have been less well studied, but review of the Eurexpress in situ hybridization database shows *Tgfβ1* expression in the medulla. Relative amplitudes of these different signaling pathways could be regulated at the level of the ligands or perhaps more likely by other intracellular signaling components or distinct signaling pathways.

In aggregate, our finding show that Smads and Wnt/β-catenin antagonistically control cell proliferation in the medullary interstitium and that the balance of these signaling pathways determines interstitial cell abundance in the postnatal kidney. Understanding this molecular crosstalk will not only contribute important concepts to the pathogenesis of CAKUT, but may also identify therapeutically tractable mechanisms that control kidney fibrosis in the adult.

## MATERIALS AND METHODS

### Mouse Strains

Animal care was in accordance with the National Research Council Guide for the Care and use of laboratory animals and protocols were approved by the Institutional Animal Care and Use Committee of Maine Medical Center. The second exon is targeted in *Smad4*^*-/+*^ and *Smad4*^*loxp/*loxp^ mice (Chu et al., 2004). Foxd1^*+/Cre*^;*Smad4*^*-/+*^ mice were crossed to *Smad4*^*loxp/loxp*^ mice to produce *Foxd1*^*+/Cre*^;*Smad4*^*+/loxp*^ (*Smad4*^*con*^) and *Foxd1*^*+/Cre*^;*Smad4*^*-/loxp*^ (*Smad4*^*IC*^) mice. *Foxd1*^*+/Cre*^;*Smad4*^*-/+*^ and *Smad4*^*loxp/loxp*^ mice were maintained on an ICR background. *R26RlacZ* mice were maintained on an FVB/NJ background.

### Reagents

FH535 (Tocris, 4344/10) was used at 15µM for *in vitro* experiments with DMEM + 2% serum as vehicle control. For in *in vivo* experiments, mice were dosed with 20mg/kg FH535 with PBS as vehicle control. Recombinant human BMP4 (100ng/ml; R&D, 314-BP), recombinant human TGFβ1 (5ng/ml; R&D, 240-B), CHIR99021 (50nM-1uM; Stegment, 04-0004), LDN-193189 (250nM; Stegment, 04-0074-02), IWR1 (10uM; Tocris, 3532/10), IWP2 (5uM; Tocris, 3533/10) and TGFβ Receptor I Inhibitor II (25nM; Tocris were also used in this study. A complete list of antibodies including source, clone name, product number and application are listed in Supplementary Table 1.

### Isolation of Primary Renal Interstitial Cells

P0 mice were individually anesthetized (16mg/kg xylazine; 105mg/kg ketamine) and perfused through the left ventricle with Dulbecco’s Phosphate-Buffered Saline (DPBS) containing magnetic particles (Spherotech AMS-40-10H) and heparin (20 units/ml) at a rate of 5ml/min with the aid of an automated syringe pump. Perfused kidneys were dissected on ice and the capsule and ureter removed. Approximately 20 whole kidneys were incubated in 3ml of Digest 1 (1.3mg/ml collagenase A; 5mg/ml pancreatin in DPBS) for 35 mins at 37°C nutating then washed three times with DPBS to remove nephrogenic zone cells. Kidneys were then dounce-homogenized and incubated in 3ml of Digest 2 (50mg/ml collagenase IV; 200 units/ml DNaseI in DPBS plus Ca^++^ and Mg^++^) for 15 mins at 37°C on a nutator. Tissue was passed through a 40-micron cell strainer with 20ml DPBS, pelleted at 300xg for 3 minutes, resuspended in 2ml DPBS, then subject to three rounds of magnetic separation to remove glomeruli. Depleted tissue was resuspended in 1ml of Digest 2, incubated for 50 mins at 37°C, then passed through a 20-micron cell separator with 10ml AutoMacs Running Buffer (ARB). Single cells were washed 2X with ARB to remove residual Digest 2 then incubated with PE-conjugated mouse anti-CD31, anti-CD45, anti-CD326, anti-Ter119 (1:10; Miltenyi) in ARB for 15 min on a nutator in the dark. Cells were washed with ARB then incubated with anti-PE microbeads (1:5; Miltenyi) for 15 min on a nutator in the dark followed by another wash with ARB. Labeled cells were subject to magnetic-activated cell sorting (MACS) and the unlabeled (negative) cellular fraction was cultured in DMEM; 10% FBS; 1% penicillin/strep.

### Immortalization and isolation of clonal lines

Isolated bulk primary renal interstitial cells were immortalized with an mCherry-tagged temperature sensitive SV40T lentivirus in which the oncogene is expressed at 33°C but not 37°C (ref). Transduced cells (MOI 50) were transferred to 33°C after 48 hours to initiate expression of SV40T. Single cells were plated by serial dilution in 96-well plates and scored daily for clonal growth. Isolated clones were cultured at 37°C for 5 days to allow cellular clearance of SV40T then screened for expression of interstitial markers (greater than a 1.5 fold change increase of transcript normalized to bulk parent population), CHIR-induced proliferation (greater than a 30% increase in EdU^+^ cells with 1uM CHIR normalized to DAPI), loss of Smad4 with tamoxifen treatment (greater than a 50% decrease in Smad4 protein with 1ug/ml tamoxifen).

### Immunoblotting

Total protein was extracted from whole kidneys as previously described (Blank et al., 2009). Immunoblotting was performed using standard procedures. Antibodies used were anti-Smad1, anti-pSmad1/5/8, anti-Smad2, anti-pSmad2, anti-Smad3, anti-pSmad3, anti-TCF4, anti-PKM1/2, anti-HisH3, anti-active (non-phosphorylated) β-catenin (1:1,000 Cell Signaling) anti-α-SMA (1:1000 Sigma), anti-PDGFRβ (1:1000 Abcam), anti-Smad4 (1:1000, Santa Cruz) and anti-β-tubulin (1:5,000, Santa Cruz). Protein levels were quantified using FIJI/Image-J software by measuring the integrated density of the indicated proteins normalized to the β-tubulin loading control.

### Quantitative RT–PCR

RNA was extracted with the RNeasy Micro Kit (Qiagen) and the concentration was measured using a NanoDrop2000 Spectrophotometer (Thermo Fisher Scientific). 1 ug total RNA was used for cDNA synthesis with qScript cDNA SuperMix (Quantabio). Quantitative RT–PCR was performed using iQ SYBR Green Supermix (BioRad). Primer sequences of genes are listed in Supplementary Table 2. Fold changes were normalized to the housekeeping gene *Gapdh* and average values (mean±s.d.) of three experimental replicates are shown. P values were calculated using a two-tailed Student’s *t*-test, and *P*<0.05 was considered significant.

### Subcellular Fractionation

Cells were treated with vehicle (DMEM) or inhibitors (25nM TGFβ RI kinase inhibitor II; 250nM LDN-193189) for 60 min followed by growth factor treatment (5ng/ml TGFβ1; 100ng/ml BMP4) for 60 min. Treated cells were transferred to fractionation buffer (20mM HEPES, 10mM KCl, 2mM MgCl_2_, 1mM EGTA, 1mM EDTA) by scraping then incubated on ice for 15 min. The cell suspension was passed through a 27-gauge needle 10 times, incubated on ice for 20 minutes, then centrifuged at 750xg for 5 minutes to produce the nuclear fraction (pellet) and the cytoplasmic fraction (supernatant). The nuclear pellet was washed with fractionation buffer by passing through a 25-gauge needle 10 times then resuspended in Tris-Buffered Saline (TBS) with 0.1% SDS. Both cytoplasmic and nuclear fraction lysates were sonicated for 3 seconds on ice at a power setting of 2-continuous before proceeding with immunoblotting.

### Immunofluorescence of monolayer cells

Cells were cultured for 5 days at 37°C then fixed in 4% paraformaldehyde (PFA) for 10 min. Cells were permeabilized with 0.5% Triton-X in PBS for 5 min, rinsed with PBS then incubated in blocking solution (PBS plus 5% serum of secondary antibody species) for 30 min. Cells were incubated in primary antibodies (diluted 1:50 in blocking solution) at 4°C overnight. Antibodies used were anti-vimentin (Sigma), anti-PDGFRβ (Abcam), anti-PDGFRα (Cell Signaling) anti-α-SMA-Cy3 (Sigma), anti-fibronectin (Santa Cruz), anti-SIX2 (Proteintech), anti-E-cadherin (BD Transduction), anti-MEIS1 (Abcam), anti-Desmin (Dako), anti-SV40T (xxx) and counterstained with Alexa Fluor 488 Phalloidin (Molecular Probes) and DAPI (Invitrogen Molecular Probes 1:10,000).

### Whole mount immunofluorescence

E14.5 and P0 kidneys were fixed in 4% PFA at room temperature for 10 or 30 minutes, respectively, then transferred to 70% ethanol at −20°C for storage. P0 kidneys were longitudinally or transversely vibratome-sectioned at 100microns directly in 70% ethanol. Whole kidneys and kidney sections were rehydrated in PBS then permeabilized with 1% Triton-X in PBS for 10 min at 4°C. Tissue was washed with PBS to remove residual detergent then incubated in blocking solution (0.01% Tween in PBS plus 5% serum of secondary antibody species) for 1 h. Lectins Dolichos Biflorus Agglutinin (DBA; Vector Laboratories) and Lotus Tetragonolobus Lectin (LTL; Vector Laboratories) were diluted 1:200 and primary antibodies anti-α-SMA-Cy3 (Sigma), anti-PDGFRβ (Abcam), anti-cytokeratin8/TROMA-1 (DSHB), anti-LEF1 (Cell Signaling), anti-BRN1 (xxx), anti-AnnexinA2 (Cell Signaling), p57kip2 (Cell Signaling), anti-CyclinD1 (Cell Signaling), anti-ki67 (Abcam), anti-TCF4 (Cell Signaling), anti-active (non-phosphorylated) β-catenin, and anti-WT1 (Santa Cruz) were diluted 1:50 in blocking solution. Tyramide Signal Amplification (TSA; PerkinElmer) was used according to the manufacturer’s protocol for whole mount immunostaining with anti-Smad4 (1:500; Santa Cruz), anti-Smad3 (1:3,000; Cell Signaling) and anti-pSmad1/5/8 (1:1000; V. Lindner) antibodies. Blocked tissue was incubated in diluted lectins/primary antibodies for 24hr at 4°C followed by three washes with blocking solution, the third wash for 24hr at 4°C. Alexa-Fluor 488/568/647 secondary antibodies (Molecular Probes) were used at 1:200 and incubated for 24hr followed by three washes with 0.01% Tween in PBS, the third wash for 24hr at 4°C. Tissue was counterstained with DAPI (1:5,000), dehydrated in ethanol, and cleared with BABB (1:1 benzyl alcohol:benzyl benzoate) before imaging with a laser scanning confocal microscope.

### In situ hybridization

The *Apcdd1* ISH probe plasmid was kindly provided by Dr. Thomas Carroll. P0 kidneys were fixed in 4% PFA overnight at 4°C, washed with PBS for 4hr, then equilibrated in 30% sucrose/PBS overnight at 4°C. After flash freezing in OCT, kidneys were cryosectioned at 20 microns and sections were fixed in 4% PFA for 10 min, rinsed three times with PBS, incubated in 20ug/ml proteinase K in PBS for 10 minutes, followed by three rinses with PBS. Sections were incubated in 1.3% triethalamine;0.375% acetic anhydride for 10 minutes, rinsed three times with PBS then incubated in hybridization buffer for 2hr. DIG-labeled riboprobes were diluted to 500ng/ml in hybridization buffer (50% formamide, 5X SSC pH 4.5, 1% SDS, 50ug/ml yeast tRNA, 50ug/ml heparin) and sections were hybridized in a humidified chamber at 68°C overnight. After two washes with 0.2X SSC for 30 min at 72°C, sections were rinsed with NTT (0.15 M NaCl, 0.1% Tween-20, 0.1M Tris-HCl pH 7.5), incubated in blocking buffer (5% heat-inactivated sheep serum, 2% blocking reagent in NTT) for 2 hours, followed by incubation in anti-DIG AP conjugated antibody (diluted 1:4,000 in blocking buffer) overnight at 4°C. After three washes with NTT for 30 min each, sections were rinsed with NTTML (0.15 M NaCl, 0.1% Tween-20, 0.1M Tris-HCl pH 9.5, 50mM MgCl_2_, 2mM levamisole) then incubated in BM purple until desired staining was reached. Stained sections were rinsed three times with PBS then mounted in glycerol for imaging.

### *In vivo* proliferation analysis

One EdU pulse (20mg/kg) was administered to P0 pups by intraperitoneal injection. After 2hr, kidneys were dissected on ice and fixed in 4% PFA for 30 min. Kidneys were vibratome sectioned and whole mount immunofluorescence staining was performed as described above. Click-it chemistry was subsequently performed on sections according to the manufacturer protocol (ThermoFisher) followed by imaging with a laser scanning confocal microscope. EdU quantification was performed with Imaris image analysis software (Bitplane). Briefly, equivalent 250×250×50 um^3^ image areas were selected in the medulla and papilla and the spot function was used to count EdU^+^ nuclei. A filter was applied to the spot function to select for PDGFRβ/α-SMA-labeled interstitial cells. Equivalent image analysis, including identical thresholding, was performed on six biological replicates (*n=6*).

### *In vitro* proliferation analysis

Cells were cultured for 5 days at 37°C followed by treatment with CHIR (50nM, 100nM, 250nM, 500nM, and 1uM) or CHIR (1uM) with FH535 (15uM) or TGFβ1 (5ng/ml) in 2% serum media for 24hr. Treated cells were pulsed with 10uM EdU for 1hr and immediately fixed in 4% PFA for 10 min. Cells were permeabilized with 0.5% Triton-X in PBS for 15 min and Click-it chemistry was performed according to the manufacturer protocol followed by counterstaining with DAPI (1:10,000). ImageJ was used to quantify EdU^+^ nuclei in three images taken from three technical replicates.

### Histology and detection of beta-gal activity

Paraffin-embedding and H&E staining of tissue was performed by the Maine Medical Center Research Institute Histomorphometry Core. For X-gal staining, whole P0 kidneys were vibratome sectioned (300uM), rinsed in X-gal buffer (5mM EGTA; 2mM MgCl_2_; 0.02% NP40; 250μM sodium deoxychalate in PBS) then fixed for 30 minutes (1% formaldehyde; 0.2% glutaraldehyde in X-gal buffer). Sections were washed 2X 10 mins with X-gal buffer then stained (5mM K_3_Fe; 5mM K_4_Fe; 0.5mg/ml X-gal in X-gal buffer) overnight at 37°C. Stained sections were washed 2X 10 mins with X-gal buffer then dehydrated through ethanol/xylenes. Dehydrated sections were embedded in paraffin then re-sectioned at 10μm and counter stained with Nuclear Fast Red.

### Single cell recombination analysis

Nephrogenic zone cells (NZCs) were isolated from E17.5 *Smad4*^*IC*^ mice as previously described (Brown et al., 2015) and labeled with PE-conjugated mouse anti-CD140a/PDGFRα (1:10 Miltenyi) followed by incubation with Anti-PE MicroBeads (1:20 Miltenyi). Labeled cells were purified by three rounds of magnetic-activated cell sorting (MACS) and single cells were manually picked from the PE-positive fraction with the aid of an EVOS digital microscope. Total DNA was amplified from single cells with the REPLI-g Single Cell Kit (Qiagen) and genotyping was performed with NovaTaq Hot Start Master Mix (Millipore) according to the manufacturer’s cycling parameters (Tm=55°C) and the primers listed in Supplementary Table 2.

### Proximity Ligation Assay

Cells were cultured for 5 days at 37°C, treated with vehicle or CHIR (1uM) and TGFβ1 (5ng/ml) for 1hr then fixed in 4% PFA for 5 min. Cells were permeabilized with 0.5% Triton-X in PBS for 5 min, rinsed with PBS then incubated in Duolink PLA blocking solution (Sigma) for 1hr at 37°C. Cells were incubated in primary antibodies against Smad3 (Cell Signaling), Smad4 (Santa Cruz), TCF4 (Cell Signaling), active (non-phosphorylated) β-catenin (Cell Signaling), active (non-phosphorylated) β-catenin (Millipore) diluted 1:50 in Duolink Antibody Diluent at 4°C overnight then washed 2X 5 min in Duolink Wash Buffer A. Cells were incubated in Duolink PLUS and MINUS probes diluted 1:5 in Antibody Diluent for 1hr at 37°C then washed 2X 5 min in Wash Buffer A. Cells were incubated with Duolink Ligase diluted 1:40 in 1X Ligation Buffer for 30 min at 37°C then washed 2X 5 min in Wash Buffer A. Cells were incubated with Duolink Polymerase diluted 1:80 in 1X Amplification Buffer overnight at 37°C then washed 2X 10 min in Duolink Wash Buffer B followed by a 1 min wash with 0.01X Wash Buffer B. Cells were counterstained with DAPI (1:10,000) and ImageJ was used to quantify the average number of foci in at least 200 nuclei from each treatment group.

### Statistical Analysis

Chi-Square testing was performed to verify that all data sets (except for Proximity Ligation Assay data) were normally distributed. Two-tailed T-tests were performed comparing normally distributed groups with p>0.05 considered significant. Proliferation analysis of FH535-treated mice and FH535-treated monolayer cells were subject to one-way ANOVA. Asterisks indicate statistical significance as follows: not significant (ns)=p>0.05; *=p<0.05; **=p<0.01; ***=p<0.00.

## Supporting information

Supplementary information for McCarthy et al

## ACKNOWLEDGEMENTS

The authors gratefully acknowledge Dr. Volkhard Lindner for providing the pSmad1/5/8 antibody and the MMCRI Histomorphometry Core for tissue processing and H&E and CD31 staining. The authors also thank Drs. Lucy Liaw, Calvin Vary, Pradeep Sathyanaryana and Ron Korstanje for their critical review of this manuscript.

