## Supplementary information for McCarthy et al for "Proliferation control of kidney interstitial cells"

**SUPPLEMENTARY TABLES & FIGURES**

**Table S1.** Primary Antibodies used in this study.

| target | company | clone | product number | application(s) |
|---|---|---|---|---|
| Troma1 | DSHB | N/A | N/A | WM |
| PdgfrB | Abcam | Y92 | Ab32570 | WM, WB, MC |
| a-SMA Cy3 | Sigma | 1A4 | C6198 | WM, MC |
| a-SMA | Sigma | 1A4 | A2547 | WB |
| Brn1 | | | | WM |
| Smad3 | CST | C67H9 | 9523 | TSA-WM, WB, PLA |
| Smad2 | CST | 86F7 | 3122 | WB |
| B-tubilin | SCB | H-235 | sc-9104 | WB |
| WT1 | SCB | | | WM |
| Smad1 | CST | N/A | 9512 | WB |
| pSmad1/5/8 | CST | N/A | 9511 | WB |
| Smad4 | SCB | B-8 | sc-7966 | TSA-WM, WB, PLA |
| Lef1 | CST | C12A5 | 2230 | WM |
| p57kip2 | CST | N/A | 2557 | WM |
| CyclinD1 | CST | 92G2 | 2978 | WM |
| Ki67 | Abcam | SP6 | Ab16667 | WM |
| active B-catenin (mouse) | Millipore | 8E7 | 05-665 | PLA |
| Tcf4 | CST | C48H11 | 2569 | WM, WB, PLA |
| active B-catenin (rabbit) | CST | D13A1 | 8814 | WM, WB, PLA |
| PKM1/2 | CST | N/A | 3186 | WB |
| HisH3 | CST | N/A | 9715 | WB |
| AnnexinA2 | CST | D11G2 | 8235 | WM |
| Desmin | Dako | D33 | M0760 | MC |
| Fibronectin | SCB | H-300 | sc9068 | MC |
| Pdgfra | CST | N/A | 3164 | MC |
| vimentin | Sigma | V9 | V6630 | MC |
| Six2 | Proteintech | | | MC |
| E-cadherin | BD | 36 | 610181 | MC |
| SV40T | | | | MC |
| Meis1 | Abcam | N/A | Ab222246 | MC |
| pSmad2 | CST | N/A | 3104 | WB |
| pSmad3 | CST | | | WB |
| pSmad1/5/8 | V. Lindner | 3131 | | TSA-WM |

WB=western blot; WM=whole mount immunofluorescence; TSA-WM=Tyramide Signal Amplification whole mount immunofluorescence; MC=immunofluorescence of monolayer cells;

PLA=Proximity Ligation Assay. Developmental Studies Hybridoma Bank (DSHB); Cell Signaling Technologies (CST); Santa Cruz Biotechnology (SCB); BD Biosciences (BD).

**Table S2.** Nucleic acid sequences of primers used in this study.

| | |
|---|---|
| **genotyping primers:** | |
| mSmad4 W4 | CTT TTA TTT TCA GAT TCA GGG GTT C |
| mSmad4 W2 | AAA ATG GGA AAA CCA ACG AG |
| mSmad4 C2 | TAC AAG TGC TAT GTC TTC AGC G |
| Cre 1 | TTC GGC TAT ACG TAA CAG GG |
| Cre 2 | TCG ATG CAA CGA GTG ATG AG |
| Hprt Cre Forward | GCT AAA GAG TTG AAC GCA AAG GTG |
| Hprt Cre Reverse | GGG CTA TGA ACT AAT GAC CCC GTA |
| Rosa26F2 | AAA GTC GCT CTG AGT TGT TAT |
| Rosa1295 | GCG AAG AGT TTG TCC TCA ACC |
| Rosa523 | GGA GCG GGA GAA ATG GAT ATG |
| **qPCR primers:** | |
| mSmad4 Forward | ACA CCA ACA AGT AAC GAT GCC |
| mSmad4 Reverse | GCA AAG GTT TCA CTT TCC CCA |
| mGapdh Forward | CATCACTGCCACCCAGAAGACTG |
| mGapdh Reverse | ATGCCAGTGAGCTTCCCGTTCAG |
| mApcdd1 Forward | CGG TGT GCT CTC ATC TAA GGT C |
| mApcdd1 Reverse | CCC ACT GAA GAC ATT GAG GAG G |
| mSnai2 Forward | TCT GTG GCA AGG CTT TCT CCA G |
| mSnai2 Reverse | TGC AGA TGT GCC CTC AGG TTT G |
| mLox Forward | CAT CGG ACT TCT TAC CAA GCC G |
| mLox Reverse | GGC ATC AAG CAG GTC ATA GTG G |
| mSmoc2 Forward | GCC AAG TGC AAA GAT CCA CAG C |
| mSmoc2 Reverse | ACA CTT GCT GGA ACT CCT TCC G |
| mAxin2 Forward | ATG GAG TCC CTC CTT ACC GCA T |
| mAxin2 Reverse | GTT CCA CAG GCG TCA TCT CCT T |
| mPdgfrb Forward | GTG GTC CTT ACC GTC ATC TCT C |
| mPdgfrb Reverse | GTG GAG TCG TAA GGC AAC TGC A |
| mLef1 Forward | ACT GTC AGG CGA CAC TTC CAT G |
| mLef1 Reverse | GTG CTC CTG TTT GAC CTG AGG T |
| mTgfb1i1 Forward | GGT CTG GAG AAT CTT CAG GAA CC |
| mTgfb1i1 Reverse | CAC CAC TGG AAG AGG AGA ATG G |

A

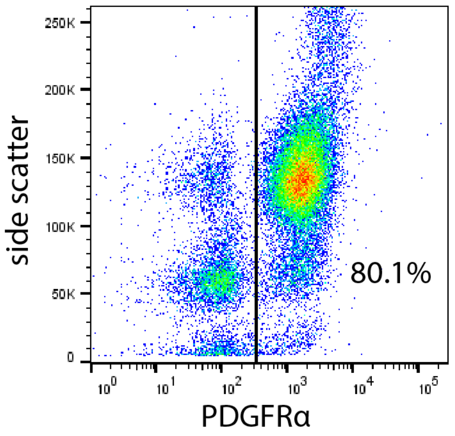


B

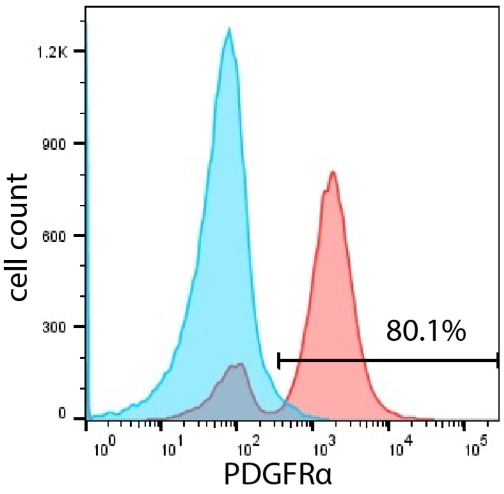


C

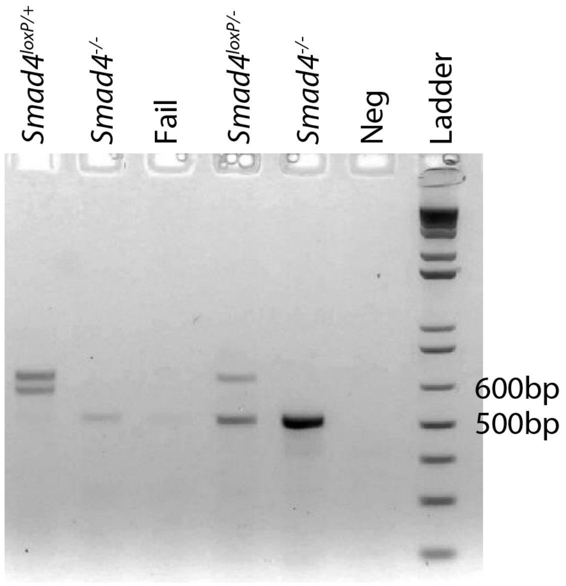


D

| | |
|---|---|
| total reactions | 74 |
| negative controls | 11 |
| *Smad4*$^{loxP/-}$ | 10 |
| *Smad4*$^{-/-}$ | 23 |
| % *Smad4*$^{loxP/-}$ recombined cells | 69.7 |
| *Smad4*$^{loxP/+}$ | 9 |
| *Smad4*$^{-/+}$ | 21 |
| % *Smad4*$^{loxP/+}$ recombined cells | 70.0 |
| total unrecombined cells | 19 |
| total recombined cells | 44 |
| % recombined cells | 69.8 |
| % PE-positive cells | 80.1 |
| % recombined PE-positive cells | 87.1 |

**Figure S1. *Smad4* is lost in renal interstitial cell precursors of *Smad4*$^{IC}$ mice.** (A) Flow cytometry showing percent PDGFRα$^{+}$ in nephrogenic zone cells enriched by 3 rounds of magnetic selection. (B) Histogram comparing the enriched cell population stained with isotype control (blue) versus PDGFRα (pink); 80.1% of cells are PDGFRα$^{+}$. (C) Gel electrophoresis of single cell genotyping PCR. Expected band sizes are 625bp for wild type, 675bp for non-recombined, and 512bp for recombined products. Amplification reactions resulting in ambiguous results (labeled “fail”) were not included in quantification. (D) Table of results for single cell genotyping.

McCarthy et al. Figure S2

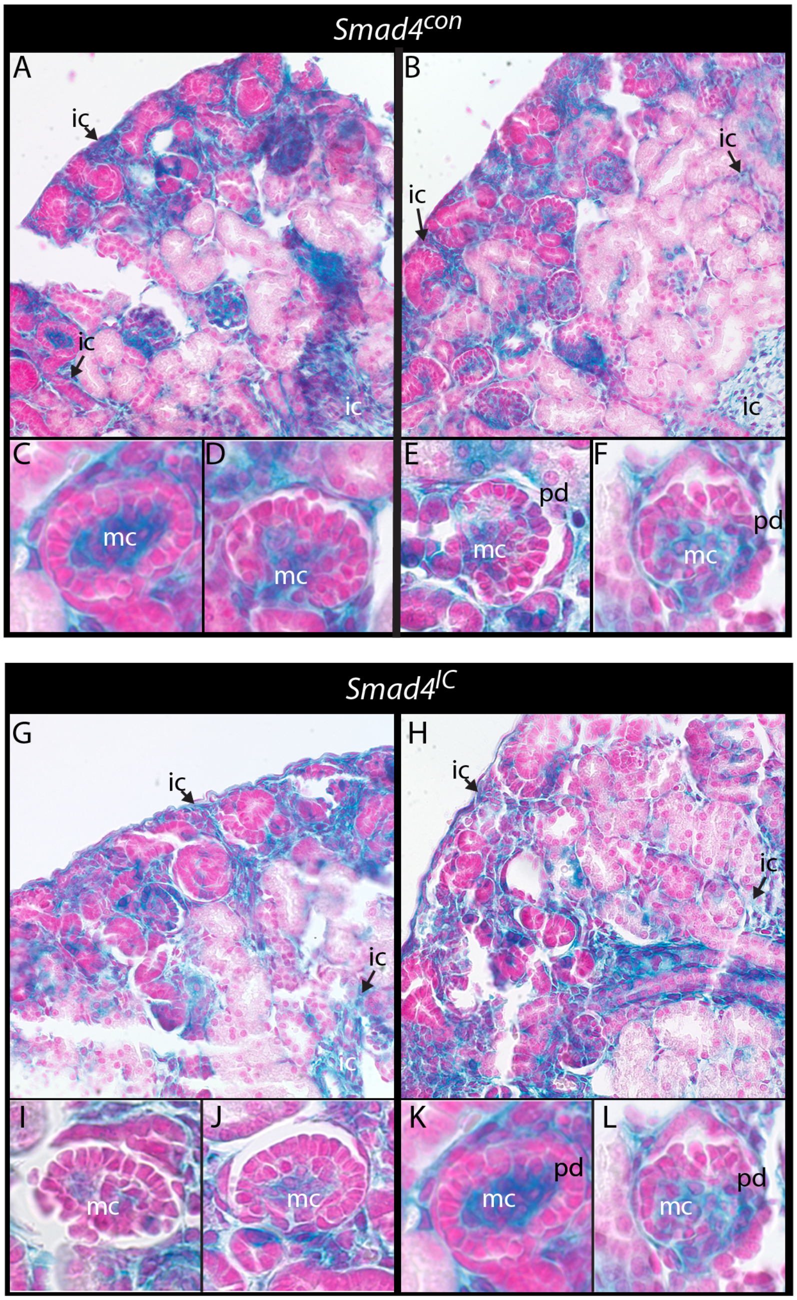


**Figure S2. Reporter gene expression reveals unchanged lineage commitment of *Foxd1*-derived cells in *Smad4*$^{IC}$.** (A, B) X-Gal staining of kidney sections from 2 *Smad4*$^{con}$ P0 pups on the R26R background. (C-F) High magnification images of glomeruli. (G, H) X-Gal staining of kidney sections from 2 *Smad4*$^{IC}$ P0 pups on the R26R background. (I-L) High magnification images of glomeruli. Abbreviations: ic, interstitial cell; mc, mesangial cell; pd, podocyte.

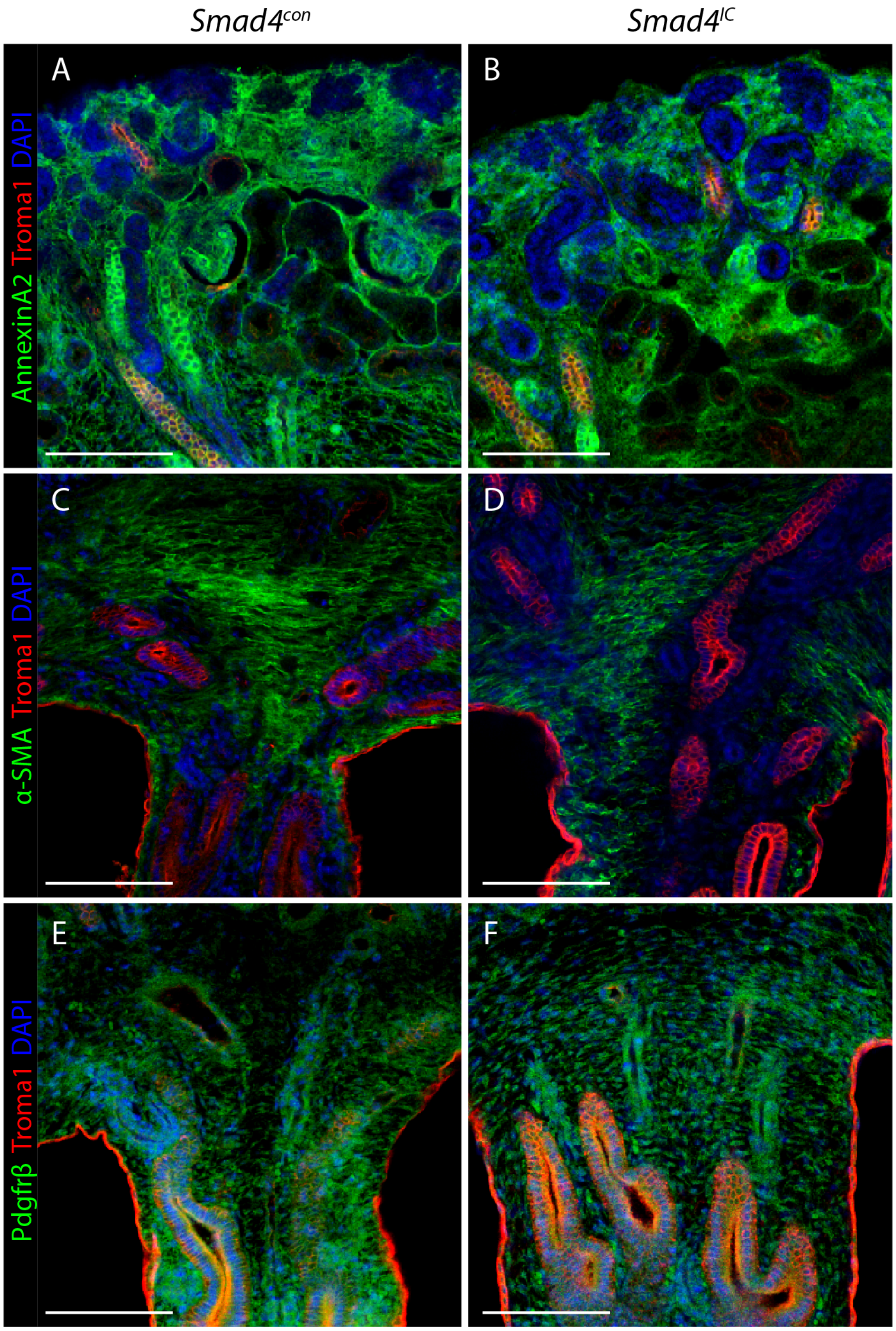


**Figure S3. No detectable phenotype is observed with loss of *Smad4* in the interstitium of E17.5 kidneys.** Cortical (A, B) and medullary (C-F) regions of E17.5 kidneys immunostained for stromal marker AnnexinA2 (A, B; representative of n=6), α-SMA (C, D; representative of n=6) or PDGFRβ (E, F; representative of n=6) and counterstained with collecting duct marker TROMA-I and DAPI. Scale bars: 200µm.

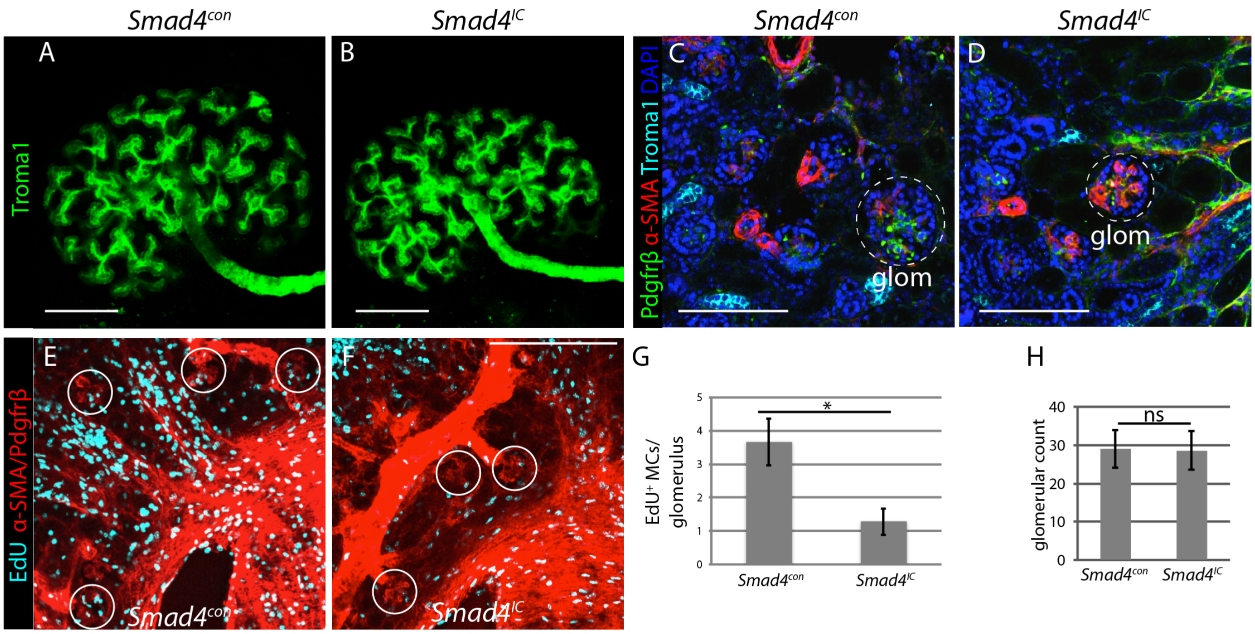


**Figure S4. Loss of *Smad4* does not affect early collecting duct branching but results in increased mesangial area and α-SMA expression and decreased mesangial proliferation.** Representative confocal images of E14.5 kidneys from *Smad4*$^{con}$ (A) and *Smad4*$^{IC}$ (B) mice immunostained with TROMA-I that were used for 3D reconstruction (n=8). (C, D) Representative sections of P0 kidneys (n=8) costained with α-SMA, PDGFRβ, TROMA-I and DAPI. Dashed circle denotes glomerulus (glom). (E, F) Representative epifluorescence images of kidney sections from EdU-treated mice with glomeruli outlined. (G) EdU+/α-SMA+ cells were quantified within each glomerulus in kidneys isolated from *Smad4*$^{con}$ and *Smad4*$^{IC}$ mice (n=6). (H) Number of glomeruli per field in *Smad4*$^{con}$ and *Smad4*$^{IC}$ kidneys at P0 (n=6). ns=p>0.05; *=p<0.05. Scale bars: 200µm in A, B.

A

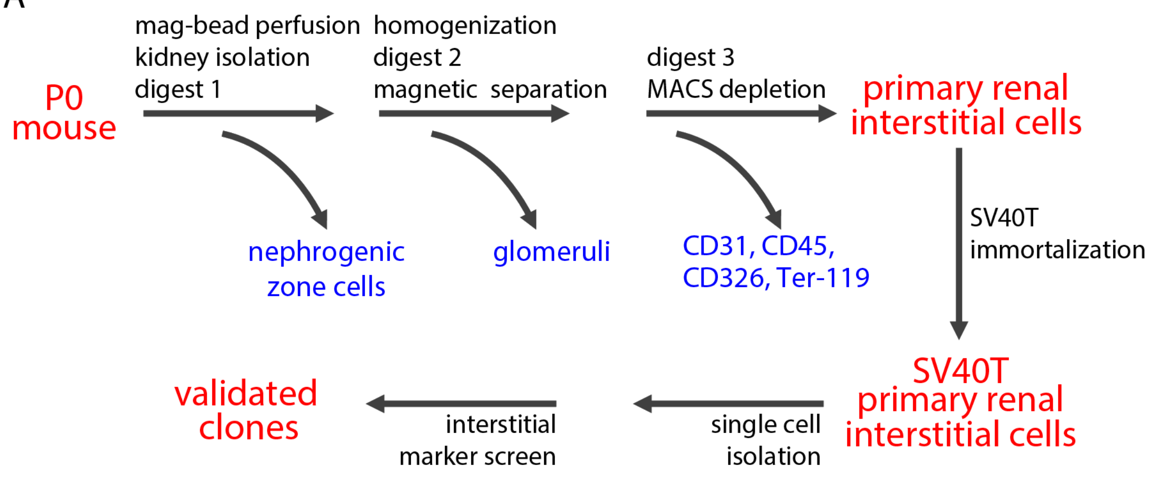


B

| | |
|---|---|
| yield | ~5x$10^6$cells/10pups |
| viability | >95% |
| PDGFRβ | 80%±2.6 |
| PDGFRα | 65%±4.7 |
| Desmin | 52%±3.0 |
| Fibronectin | 85%±5.5 |
| Vimentin | 98%±1.6 |
| Six2 | 5%±2.0 |
| DBA | <1% |
| E-Cadherin | <1% |
| GSA-B4 | <1% |

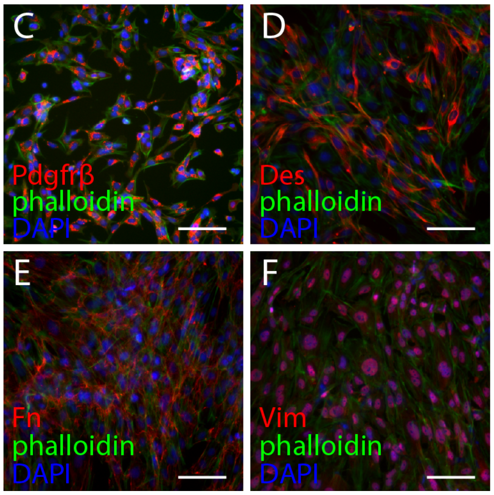


**Figure S5. Validation of primary renal interstitial cell (PRIC) isolation method.** (A) Schematic of method utilized to generate PRIC immortalized clones. (B) Table of yield, viability and quantification of marker analysis of monolayer PRICs 24 hours post isolation. Monolayer PRICs immunostained with PDGFRβ (C), desmin (D), fibronectin (E) or vimentin (F) counterstained with Alexa Fluor 488 phalloidin to label filamentous actin. Scale bars: 10µm.

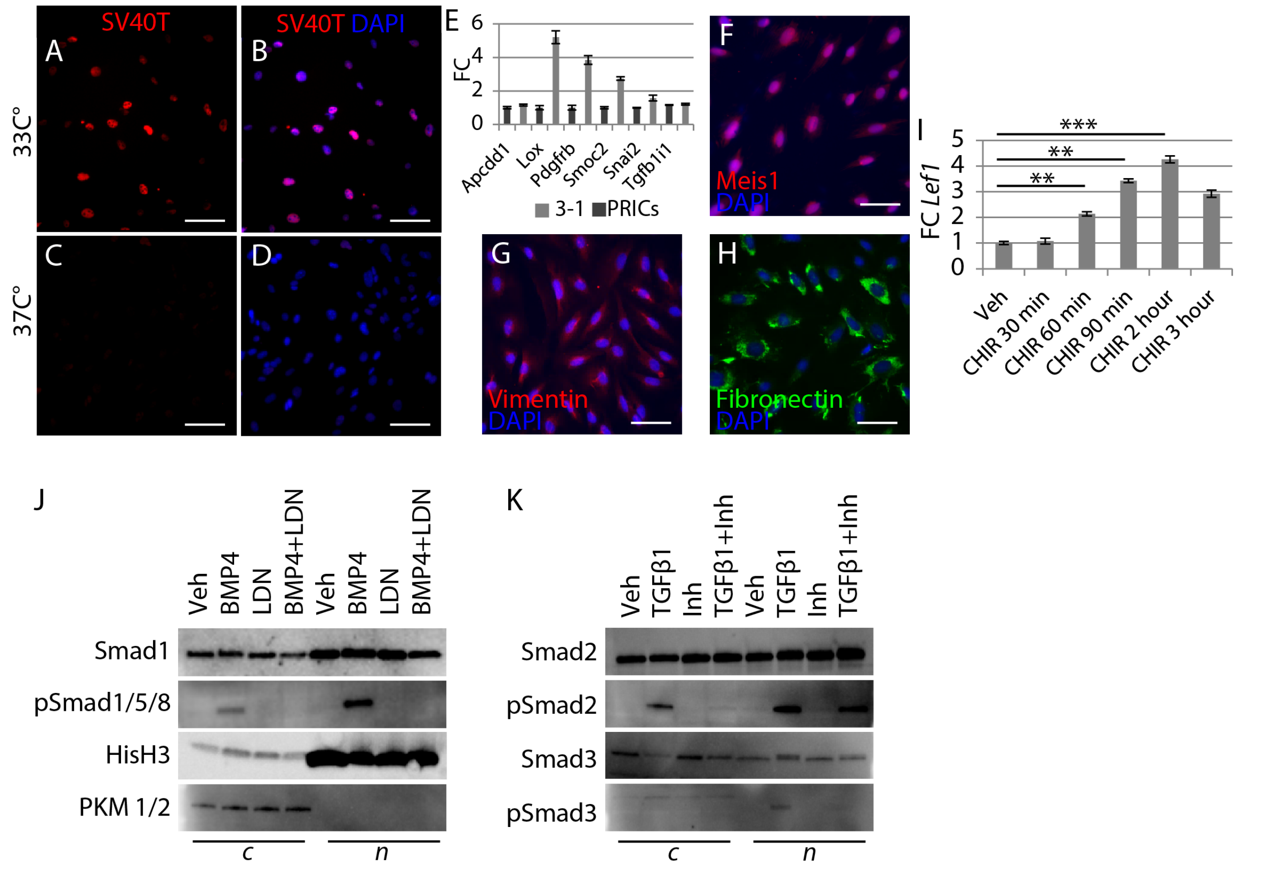


**Figure S6. Clone 3-1 maintains interstitial marker expression and responds to stimulation of the Wnt, BMP and TGFβ pathways.** SV40T immunostaining with DAPI costain of monolayer 3-1 cells grown at 33°C (A, B) or 37°C for 5 days (C, D). (E) RT-QPCR of an interstitial cell marker gene panel on cDNA from 3-1 cells and the primary renal interstitial cell (PRIC) population from which they were immortalized. Immunostaining for MEIS1 (F), vimentin (G) or fibronectin (H) of monolayer 3-1 cells grown at 37°C for 5 days. (I) Time course of *Lef1* transcript levels after treatment with 1μM CHIR normalized to vehicle-treated. (J) Immunoblot for Smad1 and its activated form pSmad1/5/8 in cytoplasmic (c) or nuclear (n) fractions of 3-1 cells stimulated for 1 hour with BMP4 (100ng/ml) with or without the LDN193189 BMP-Smad inhibitor. (K) Immunoblot for Smad2, its activated form pSmad2, and Smad3 in cytoplasmic (c) or nuclear (n) fractions of 3-1 cells stimulated for one hour with TGFβ1 (5ng/ml) with or without TGFβ Receptor I Inhibitor II. *=p<0.05; **=p<0.01; ***=p<0.001. Scale bars: 10μm in A-H.

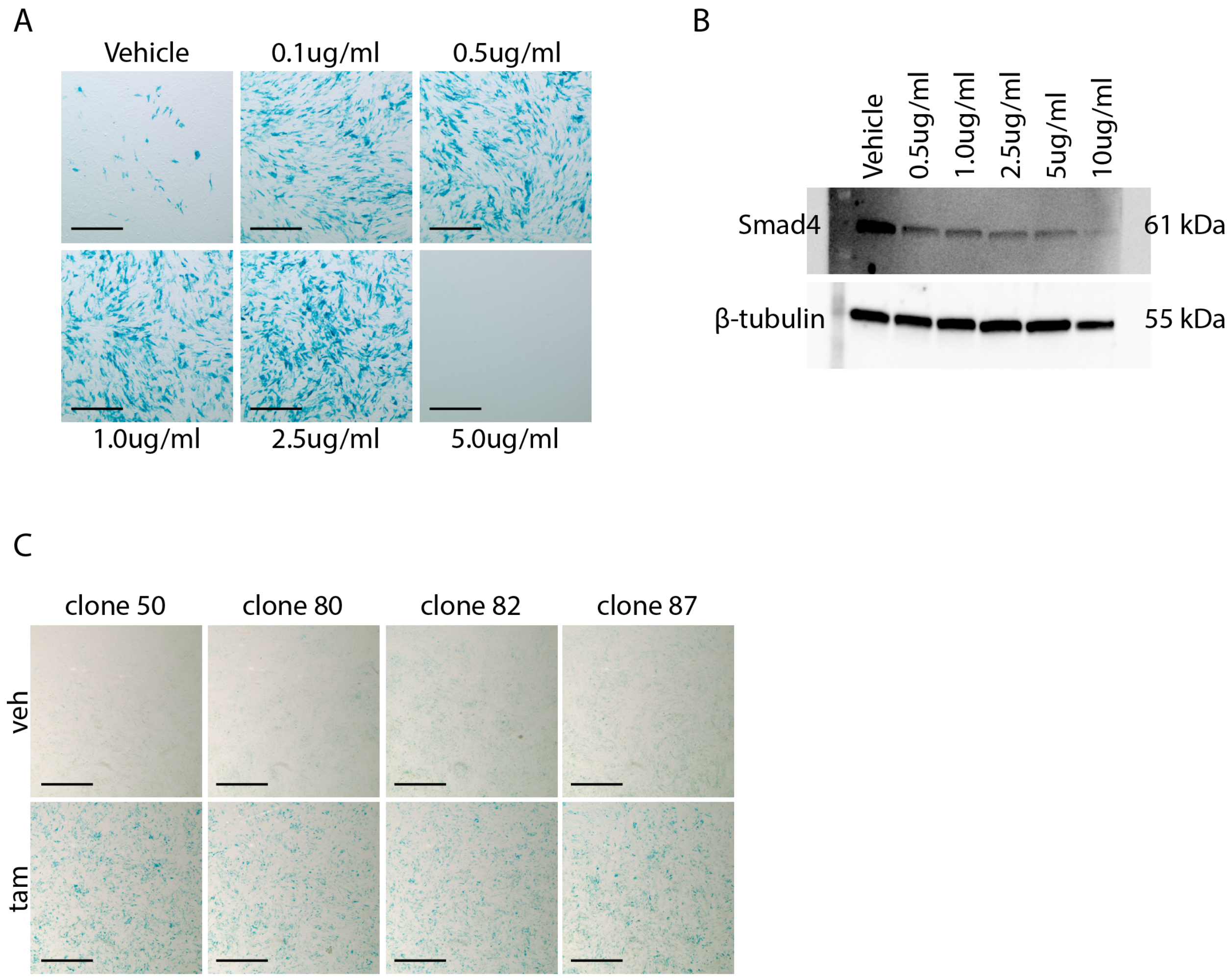


**Figure S7.** X-Gal staining (A) and immunoblot for Smad4 (B) of monolayer *Hprt*$^{Cre}$*;Smad4;R26R* immortalized cells treated with increasing tamoxifen concentrations for 24 hrs. β-tubulin was used as a loading control. (C) X-Gal staining of monolayer *Hprt*$^{Cre}$*;Smad4;R26R* clones 50, 80, 82, 87 treated with vehicle or 1µg/ml tamoxifen for 24 hrs. Scale bars: 20µm.
